# Mapping the North American Terrestrial Carbon Cycle: A Process-based Reanalysis Using State Data Assimilation (SDA)

**DOI:** 10.64898/2026.02.25.708030

**Authors:** Dongchen Zhang, Jonathan Huggins, Qianyu Li, Shashank Ramachandran, Shawn Serbin, Cameron Webb, Zhenpeng Zuo, Michael Dietze

## Abstract

The ability to accurately assess ecosystem C budgets across scales from individual sites to continents is essential for C accounting, management, and ultimately mitigating climate change. State data assimilation (SDA) provides a framework for harmonizing observations with models, while robustly accounting for and reducing multiple sources of uncertainty. In this study, we employed a hybrid SDA framework that combines process-based terrestrial biosphere modeling, hierarchical Bayesian inference, and machine learning to harmonize bottom-up and remotely-sensed data streams for 8,000 pre-selected 1km^2^ locations across North America within a hybrid structure. Combining bottom-up soils data (SoilGrids) with spectral (MODIS and Landsat) and microwave (SMAP) remote sensing helps constrain the major C and water stocks through space and time. Machine learning is used both to identify and correct systematic errors in the process model (SIPNET) and to interpolate the pre-selected locations onto a 1km grid, making it computationally feasible to generate annual ensemble maps of the North American carbon budget. Furthermore, the uncertainties for each variable were reduced compared to those from observations or models alone. Spatiotemporal analysis showed a slight decrease in aboveground biomass (AGB) across the western US, a loss of leaf area across the boreal, and a slight greening of the Alaskan tundra. The uncertainty trends suggest a significant reduction in the uncertainty about soil organic carbon (SOC), the largest C reservoir. Validation results show that we accurately estimate C pools, compared to the assimilated data streams and held-out observations of AGB from GEDI, ICESat-2, and the US FIA, and SOC from the ISCN network. Our ML-debiasing algorithm further improved the accuracy of major C pools (AGB, SOC). In general, our continental SDA framework will facilitate global C MRV (monitoring, reporting, and verification) by providing accurate and precise C-cycle estimates, along with their corresponding spatiotemporal uncertainties.

## 1. Introduction

Carbon emissions are driving global climate change, contributing to global warming, sea-level rise, an increase in extreme weather events, and other societal impacts (Matthews et al., 2009; Amirkhani et al., 2022). At the same time, carbon sinks in the land, ocean, and atmosphere have also increased, with terrestrial ecosystems capturing about one-third of the total carbon emissions (Friedlingstein et al., 2023). However, terrestrial sinks are also the largest source of uncertainty in the global C budget, are highly variable in space and time, and can be strongly affected by anthropogenic activities and natural disturbances (Arora & Boer, 2010; Harris et al., 2016). This, in turn, creates a pressing need for improved monitoring, reporting, and verification (MRV).

Despite efforts to accurately quantify terrestrial carbon budgets, current land sink estimates remain uncertain due to inconsistencies in space, time, and standard of field measurements (Acharya et al., 2022), discrepancies between field measurements and theory (Raupach et al., 2005), variations among models (Padrón et al., 2022), and the impact of human activities amid global climate and environmental changes. These challenges make it difficult to obtain accurate and economic C MRV. Therefore, it is crucial to develop methods that reduce uncertainty and enable evaluation of the latest trends in land carbon sinks across the continent.

At the same time, significant progress has been made in reducing uncertainties in land carbon estimates by combining field measurements, remote sensing, and terrestrial biosphere models (TBMs). While field-based networks for direct measurements of carbon stocks (e.g., national forest inventories, International Soil Carbon Network (ISCN), Nave et al., 2016) and fluxes (e.g., FluxNET, Baldocchi et al., 2001) are critical for anchoring C budgets and groundtruthing indirect carbon estimates (e.g., remote sensing, TBMs), ongoing fieldwork is labor-intensive and costly. Additionally, field measurements are often challenging to scale directly to wall-to-wall estimates (Williams et al., 2005). Remote sensing, however, offers scalable and comparable gridded estimates of aboveground biomass (e.g., LandTrendr, Hooper & Kennedy, 2018; GEDI, Dubayah et al., 2022; ICESat-2, Duncanson et al., 2023), LAI (MODIS, VIIRS, Myneni et al., 2015), SM (SMAP, Reichle et al., 2023), and CO2 (OCO-2, Eldering et al., 2017). Nonetheless, both field-based and remote sensing estimates typically measure a single carbon pool or flux at a time, necessitating the integration of these measurements to build a comprehensive picture of how ecosystem C budgets vary across space and time.

By contrast, TBMs are mechanistic representations of ecosystems based on current understanding and hypotheses of ecological processes. Their ability to simulate the carbon, water, and energy flows of the biosphere (Yuan et al., 2024) thus provides a flexible platform for combining information from various data sources. TBMs are generally modularized into separate components that represent key processes (e.g., photosynthesis, Farquhar et al., 1980; carbon allocation, Litton et al., 2007; respiration, Richardson et al., 2006; turnover, and decomposition, Berg et al., 2000). Compared with direct field measurements, TBMs are easier to scale due to their modular design and consistent spatiotemporal scales across modules. Moreover, TBMs also have minimal data requirements (Wesselkamp et al., 2024), making them applicable to data-sparse regions (e.g., tundra, tropics). Because of this, TBMs are crucial in regional to global land carbon projects, such as CMIP (Coupled Model Intercomparison Project, Dunne et al., 2024), TRENDY (Trends and drivers of the regional scale terrestrial sources and sinks of carbon dioxide, Sitch et al., 2024), and RECCAP2 (the second phase of the REgional Carbon Cycle Assessment and Processes Project, Ciais et al., 2020).

However, due to the limitations in our understanding of land biosphere processes, different assumptions and process representations across TBMs (Rogers et al., 2017), and the substantial uncertainties in model calibrations (Dokoohaki et al., 2021), TBMs exhibit significant uncertainties in simulated carbon cycling (Friedlingstein et al., 2023; O’Sullivan et al., 2022) and often fail to match ground-based measurements accurately across C cycling estimates (e.g., SOC, Todd-Brown et al., 2013; GPP, Schaefer et al., 2012). Furthermore, TBMs applied to regional to global studies have relatively coarse spatial resolutions (e.g., 0.5 - 1° resolution for TRENDY, 1 - 2.5° for CMIP6, 1° for RECCAP2), making it challenging for the accuracy targeting and management of C MRV. Ultimately, models by themselves are not a record of what actually occurred in an ecosystem. Therefore, it is essential to combine models with data to minimize the bias and uncertainties associated with TBMs.

**Table 1.** Naming conventions and units.

| Variable full name | Abbreviation | Unit |
| --- | --- | --- |
| Aboveground Biomass | AGB | Kg/m2 |
| Leaf Area Index | LAI | m<sup>2</sup>/m<sup>2</sup> |
| Soil Moisture | SM | % Volume |
| Soil Organic Carbon | SOC | Kg/m2 |

State Data Assimilation (SDA) provides the solution by constraining key components of modeled processes using data. Specifically, SDA is a statistical method that can integrate multiple observations with key model state variables (e.g., carbon pools), assuming that neither TBMs nor observations can precisely represent the entire ecosystem (Williams et al., 2005). Furthermore, SDA offers the ability to obtain continuous estimates across space, time, and ecological processes provided by TBMs, indirectly constraining carbon states or processes that are difficult to measure directly in space or time (Dietze, 2017), thereby reducing uncertainty in carbon budget estimates (Zhang et al., 2025). Hence, SDA offers a potential solution by combining informative direct observations with a comprehensive mechanistic understanding provided by TBMs, while also enabling comprehensive uncertainty propagation (Li et al., 2024). Because of this, SDA has been increasingly used across TBMs, especially the critical quantities in carbon cycle studies, such as LAI (Ramos et al., 2018; Kumar et al., 2019; Li et al., 2019; Chen & Tao, 2020) and SM (Sawada et al., 2015; Fairbairn et al., 2017), as well as other combinations (e.g., LAI + AGB, Dokoohaki et al., 2022 & Raczka et al., 2021; LAI + fraction of absorbed photosynthetically active radiation (faPAR), Viskari et al., 2015).

Although applying SDA reduces bias and uncertainty about the C cycle, systematic patterns in residual errors may still exist that elevate uncertainty or induce compensating errors (Scherrer et al., 2023). Therefore, SDA workflows increasingly incorporate bias-correction algorithms that iteratively adjust the estimated model simulations based on statistical relationships between residual errors and other covariates at the previous time step (Weir et al., 2021; Qin et al., 2022). Such efforts are part of a broader trend toward hybrid systems that combine the speed and flexibility of machine learning (ML) methods with physical realism and interpretability of TBMs (Wesselkamp et al., 2024). Another example of a hybrid approach is the use of ML emulators, which serve as efficient surrogates for computationally demanding TBMs and/or SDA workflows (Houborg et al., 2018; Rani et al., 2022; Padarian et al., 2020).

To generate additional projections beyond the training dataset (e.g., spatial interpolation), and to achieve accurate, continuous, and up-to-date C MRV, as well as interpretable ML outputs, we developed a continental-scale hybrid SDA system that combines the SIPNET (Simplified Photosynthesis and Evapotranspiration) model with an unprecedented number of C observations. By combining SDA with ML emulation and downscaling, our system can produce an ensemble of gridded estimates of continental-scale carbon and water budgets that captures their uncertainties. We apply this system to North America (NA) using data from 2012 to 2024 at a 1-km resolution to answer the following questions about the key factors that impact the accuracy and uncertainty of C MRV:

**Q1.** What are the general spatial-temporal patterns and trends of C budgets and the associated uncertainties, and their spatial-temporal correlations to explanatory covariates?

**Q2.** What are the patterns, across space, time, and C pools, of the residual errors in the hybrid SDA when compared to the corresponding data constraints and other data products during the validation?

**Q3.** Can the ML bias-correction improve the accuracy of our predictions? What factors are associated with model biases across space, time, and C pool?

## 2. Data and methods

To manage computational costs, avoid redundant estimates, and account for spatial autocorrelation, the first step in our SDA workflow is to select representative locations (Sec 2.1). Locations were selected using a stratified random approach based on a cluster analysis of 17 eco-climatic attributes for each land cover class. Second, for each location, we extracted a variety of information of model inputs and data constraints within the SDA (Sec 2.2). Third, we ran the SDA for the selected sites (Sec 2.3). Fourth, we developed a ML emulator for the SDA outputs (Sec 2.4). Finally, we performed a series of analyses to answer the three key questions raised in introduction (Sec 2.5). The overall workflow was implemented within the Predictive Ecosystem Analyzer (PEcAn, https://github.com/PecanProject/pecan, version 1.10.0), an open-source community platform for land model-data integration (Dokoohaki et al., 2022; Fer et al., 2018; LeBauer et al., 2013; Raiho et al., 2020). Within PEcAn, the SDA is implemented using NIMBLE v0.12.1 (de Valpine et al., 2017) while the rest of PEcAn is implemented in R (R version 4.1.2, http://www.r-project.org) unless otherwise noted.

### 2.1. Site Selection and Study Area

#### 2.1.1. Study Area

This study investigates the North American (NA) continent (-179° W to -20° W; 7° N to 85° N) at an ∼1 km spatial resolution (MODIS Sinusoidal tile grid) on July 15th each year from 2012 to 2024, focusing on carbon cycling during the peak growing season across most of the continent. It covers over 21 million km² of vegetated land, including Siberia, Central America, the Caribbean, Mexico, the USA, and Canada. Using the site-selection workflow described below, we selected 8,000 locations representing eight land cover classes (Figure 1) based on the MODIS Land Cover classifications (Friedl et al., 2022).

**Figure 1.**
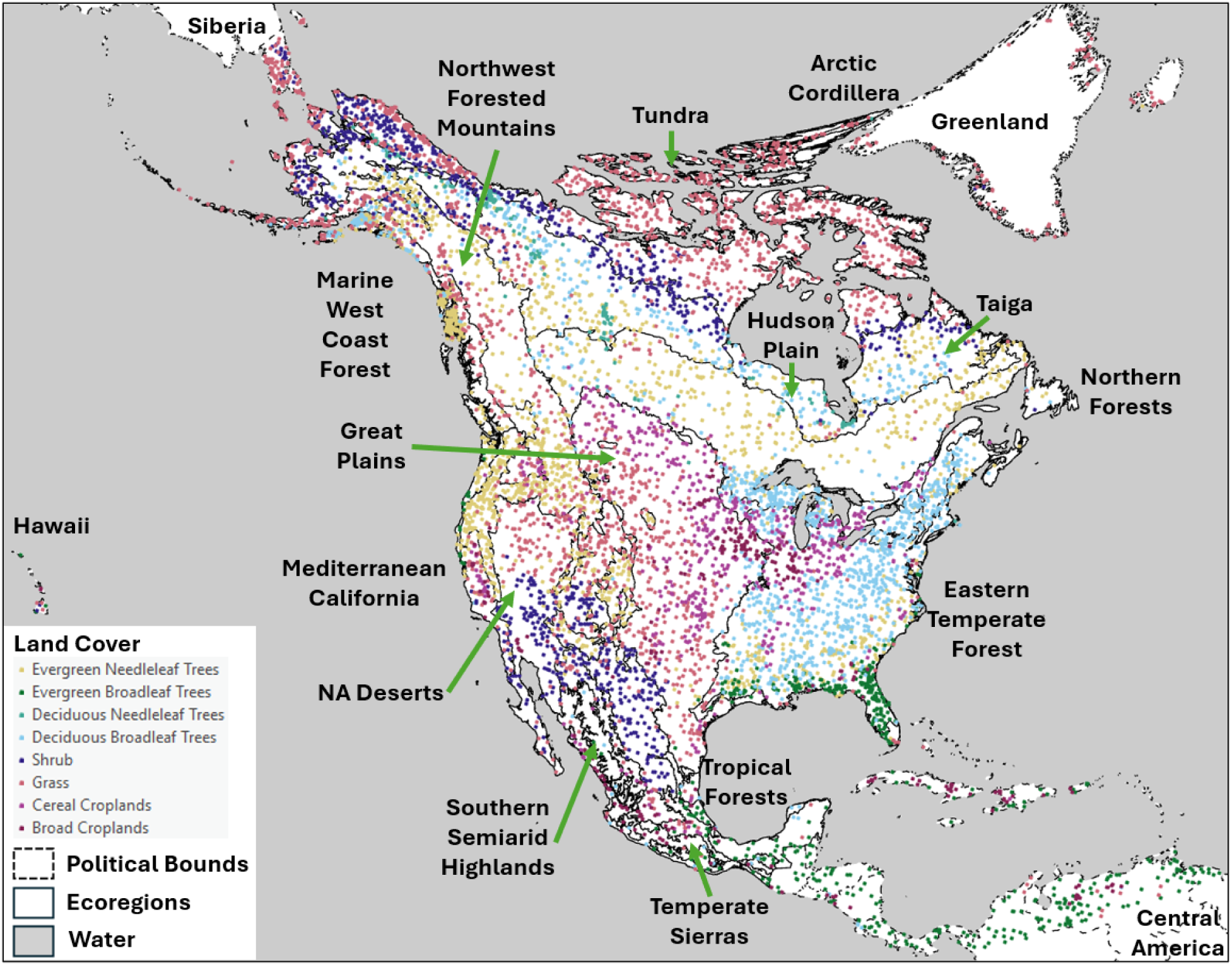
Pre-selected 8,000 locations based on our site-selection algorithm. The left table shows the abbreviations of ecoregions labeled in this figure. The map is in a CONUS Albers projection (EPSG: 4269) to preserve area across latitude.

#### 2.1.2. Site Selection

The site-selection algorithm was developed to select representative locations across eco-climatic variations, conditional on each vegetation type from the MODIS PFT land cover classifications. In this step (Figure 2), we first compiled the locations of ∼400 “anchor” sites, defined as locations with ground-based networks (e.g., Ameriflux, MexFlux, NEON, LTER, LTAR, ForestGEO, CZO, Phenocam) that are of scientific interest, and combined this with the ∼500 sites from our previous CONUS SDA (Dokoohaki et al., 2022), which were selected using an algorithm analogous to that described below.

**Figure 2.**
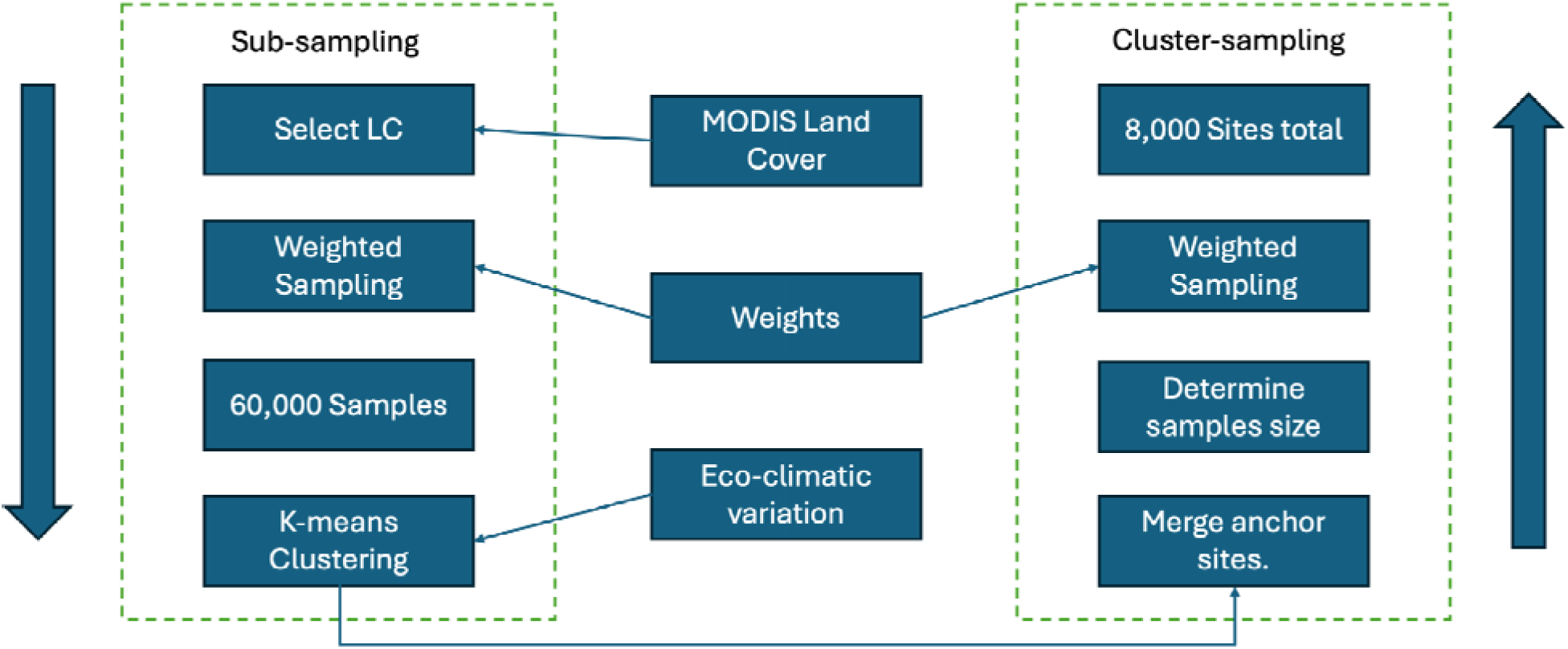
Site selection workflow. We first generate 60,000 random subsamples per MODIS land cover type using weighted sampling, prioritizing FIA and GEDI coverages. Next, we generate k-means clusters based on the eco-climatic variables (Table 2), exclude the pre-selected sites (e.g., NEON, Ameriflux, FluxNet), and determine the optimal cluster size by the elbow location in the WCSS curve. Finally, weighted sampling is applied to each cluster.

**Table 2.** Environmental covariates used for k-means clustering, debiasing, and ML emulators. ^1^Forest age 2010 base map updated annually using MODIS land cover and extended to non-forest pixels.

| Data Source | Temporal Domain | Spatial Resolution | Temporal Frequency | DOI |
| --- | --- | --- | --- | --- |
| ERA5:<br>temperature,<br>precipitation,<br>solar radiation,<br>vapor pressure | 2012-2024 | 30 km | annual | 10.24381/cds.ad<br>bb2d47 |
| STRM DEM | 2000 | 90 m | static | [https://srtm.cs<br>i.cgiar.org](https://srtm.cs<br/>i.cgiar.org). |
| Forest Age | 2010 | 1 km | annualized<sup>1</sup> | [https://doi.org/1<br>0.17871/Forest<br>AgeBGI.2021](https://doi.org/1<br/>0.17871/Forest<br/>AgeBGI.2021) |
| GEDI level 4B:<br>number of<br>samples | 2019-04-18 to<br>2023-03-16 | 1 km | static | 10.3334/ORNLD<br>AAC/2299 |
| MODIS<br>MCD12Q1 Land<br>Cover | 2001 to 2024 | 500 m | annual | 10.5067/MODIS/<br>MCD12Q1.061 |
| AGB 2010 | 2010 | 300 m | static | 10.3334/ORNLD<br>AAC/1763 |
| SoilGrids: ORC,<br>pH, nitrogen,<br>and sand | NA | 250 m | static | [https://doi.org/10<br>.5194/soil-7-217-2021](https://doi.org/10<br/>.5194/soil-7-217-<br/>2021) |
| FIA Plot Density | 2000 to 2024 | NA | static | [https://doi.org/10<br>.2737/RDS-2001-FIADB](https://doi.org/10<br/>.2737/RDS-<br/>2001-FIADB) |

Next, we calculated the number of locations for each land cover class proportionally to its spatial coverage, with a minimum of 200 sites per class (Figure 2), totaling 8,000 sites, to balance computational demand and representation of eco-climatic gradients. The remaining locations, excluding the anchor sites, were weighted for sampling to aid in later FIA and GEDI validations (Sec 2.5.2). The weighting was conducted under the CONUS Albers projection, which preserves area across latitudes, and the weight layer was calculated as the number of by-pixel FIA plots and high-quality GEDI waveforms (Table 2). Additionally, because the GEDI orbit stops at 52°N, samples for each land cover class were further split along this latitude proportionally to area.

We then assembled a suite of environmental covariates selected to explain variability in carbon pools and fluxes (Table 2). A global forest age map circa 2010 (Besnard et al., 2021) was used to represent the time elapsed since the most recent disturbance, adjusted to 2024, and supplemented with the MODIS land cover product to account for post-2010 land cover changes and the age of non-forest land cover classes. Additional covariates include the global AGB in 2010, annual median maps for MODIS LAI and SMAP root zone SM, elevation from the STRM digital elevation model dataset (Jarvis et al., 2008), and soil properties (e.g., organic carbon content (ORC), pH, nitrogen, and sand proportion) from the SoilGrids database (Hengl et al., 2017). Climatic covariates, including temperature, precipitation, solar radiation, and vapor pressure, were generated based on ERA5 annual mean estimates. For clustering, all data layers were normalized to a 0–1 scale based on their min/max values, except land cover, which was converted to a factor.

Then, we employed the Hierarchical k-means clustering method (Alboukadel & Fabian, 2020) to identify unique clusters for each land cover class, using 60,000 weighted sampled locations per class to accommodate the memory limitations of the algorithm. The k-means clustering was executed based on the maximum total within-cluster sum of squares (WCSS) obtained based on the elbow point on a plot of WCSS versus cluster sizes from 1 to 20. After dividing the number of locations to be sampled evenly among clusters, we removed the anchor sites and sampled from each cluster in proportion to the FIA and GEDI weight layer.

### 2.2 Data and Data Preprocessing

#### 2.2.1. Initial Conditions (ICs)

The initial conditions (ICs) define the starting states of the ecological model (Sec 2.3.1). To initialize LAI, we used the maximum MODIS LAI observations from 2011-06-01 to 2011-08-31 after removing outliers (>0.95 or <0.05 quantiles), with QA/QC values of either “000” or “001”. To avoid singularity issues in Bayesian analyses, we set a minimum of 0.66 standard deviation (SD) for LAI (Viskari et al., 2015). The AGB initial conditions were based on a 2010 global AGB map (Spawn et al., 2020). SM was initialized from meteorological reanalysis (Dorigo et al., 2019). We eventually generated 100 ensembles for each IC and location using the reported mean and SD for each IC. All ICs were sampled using the zero-truncated normal sampling (Olaf et al., 2018) independently for each location.

SOC was initialized using the ISCN database (Nave et al., 2016) with less than 200 cm of soil depth. To reduce the potential double use of data in both initialization and SDA between ISCN and SoilGrids, SOC priors were generated by sampling ISCN measurements with respect to each of the fifteen EPA level 1 ecoregions (Omernik, 1995). The global mean and SD will be used for places outside of the EPA map.

**Table 3.** Data used as model inputs. Here, we list the data source, temporal domain, and frequency, as well as the spatial resolution and data DOI for model drivers (ERA5 meteorology, MODIS phenology), initial conditions (e.g., AGB, LAI, SM, and SOC), and soil physical variables.

| Data Source | Temporal Domain | Spatial Resolution | Temporal Frequency | DOI |
| --- | --- | --- | --- | --- |
| ERA5 | 2012 to 2024 | 30 km | 3-hour | 10.24381/cds.adbb2d47 |
| MODIS Phenology | 2012 to 2023 | 500 m | 1-year | 10.5067/MODIS/MCD12Q2.061 |
| AGB 2010 | 2010 | 300 m | NA | 10.3334/ORNLD AAC/1763 |
| MODIS LAI | 2011 | 500 m | 8-day | 10.5067/MODIS/MOD15A3H.006 |
| Gridded Soil Moisture | 2011 | 0.25 degree | 10-day | 10.24381/cds.d7782f18 |
| ISCN SOC | 2012 to 2014 | NA | NA | 10.17040/ISCN/1305039 |
| SoilGrids soil texture data | NA | 250 m | static | 10.5194/soil-7-217-2021 |

#### 2.2.2. Model Inputs

Model parameter ensembles were sampled from by-PFT joint posterior parameter distributions generated through previous calibrations using a Hierarchical Parameter Data Assimilation (HPDA) approach against carbon and water flux measurements from 22 Ameriflux towers (Dokoohaki et al., 2022; Fer et al., 2018, 2021). Phenological drivers, including leaf-on and leaf-off dates, were extracted from the MODIS Land Cover Dynamics (Friedl et al., 2022) database, which provide annual global land surface phenology metrics from 2001 to the present at a 500 m spatial resolution. We used the “mid-green-up” and “mid-green-down” bands when EVI2 (enhanced vegetation index 2) crossed 50% of the seasonal amplitude to identify leaf-on and leaf-off dates. Ensembles of site-specific soil texture parameters (e.g., soil water holding capacity & drainage rate) were generated based on the SoilGrids dataset (Supplement 1) using the look-up tables and pedotransfer functions in Cosby et al. (1984). The ERA5 reanalysis was used as meteorological drivers (Hersbach et al., 2023) given its continuous global coverage with ten ensembles, which were randomly assigned to the 100 model ensembles.

#### 2.2.3 Observational Data Constraints

This study simultaneously assimilated AGB, LAI, SM, and SOC (Table 2), building on our previous work assimilating the same constraints across CONUS NEON sites (Zhang et al, 2025). We used LandTrendr product (Hooper & Kennedy, 2018) as the AGB constraint, which utilizes the LandTrendr estimates of the Landsat composite within CONUS, calibrated against forest inventory data, which provides AGB mean and uncertainty estimates until 2017, and only AGB means from 2018 to 2023. To predict uncertainties for 2018 to 2023 we trained the RandomForest algorithm against the reported uncertainties from 2012 to 2017 using covariates from the site-selection algorithm (Supplement 2).

MODIS LAI observations were extracted from June 1st to August 31st for each year (with the QA/QC values fall within “000” or “001”) and the 0.025-0.975 quantiles to avoid noisy estimates due to cloud cover. We then selected the estimates that are closest to July 15 for each year and location, with 90% of the values falling within the 8 days of the target date (see Figure S1). Similar to LAI IC, we assigned a minimum SD of 0.66 to the extracted LAI observations.

The SM is constrained by the SMAP Level 4 root-zone SM estimates, which use a data assimilation system to merge L-band brightness temperature observations with the NASA Catchment land surface model (Reichle et al., 2023). The estimates provide continuous SM to 100 cm depth (the same root-zone definition as SIPNET) with associated uncertainties. In this study, we extracted SMAP estimates at the mid-night of July 15 each year from 2015 to 2024.

SOC estimates were obtained to a depth of 200cm from the OCD band of SoilGrids data (relationship between ORC and OCD bands can be found in suplement 3). Since uncertainty estimates are given as a mean and three quantiles (0.05, 0.5, 0.95) for each soil layer, we converted these into an overall column-integrated mean and standard deviation by fitting Gamma distributions across six soil depths (0 - 5cm, 5 - 15cm, 15 - 30cm, 30 - 60cm, 60 - 100cm, and 100 - 200cm) and aggregating the distribution’s mean and variance through the entire soil profile.

**Table 4.** Details for the data used as observations. Here, we summarized the data details by their name, temporal domain, frequency, spatial resolution, and the data DOI.

| Data Source | Temporal Domain | Spatial Resolution | Temporal Frequency | DOI |
| --- | --- | --- | --- | --- |
| LandTrendr AGB | 2012 to 2023 | 30 m | 1-year | [10.1088/1748-9326/aa9d9e](https://doi.org/10.1088/1748-9326/aa9d9e) |
| MODIS LAI | 2012 to 2024 | 500 m | 8-day | http://doi.org/10.5067/MODIS/MOD15A3H.006 |
| SMAP level 4 root zone soil moisture | 2015 to 2024 | 9 km | 3-hour | [doi:10.5067/02LGW4DGJYRX](https://doi.org/10.5067/02LGW4DGJYRX) |
| SoilGrids OCD | NA | 250 m | NA | https://doi.org/10.5194/soil-7-217-2021 |

### 2.3. SDA workflow

In this study, we employed the SDA approach, the Tobit Gamma Ensemble Filter (TGEnF), a variant of the Tobit Wishart Ensemble Filter (TWEnF) (Raiho et al., 2020; Dokoohaki et al., 2022), to statistically harmonize observations and model forecasts. Similar to other ensemble-based assimilation approaches (e.g., the Ensemble Kalman Filter (Evensen, 2003)), this approach begins by generating an ensemble prediction of the modeled pools and fluxes, adds model process error, and then uses this ensemble mean and covariance to parameterize a multivariate Bayesian prior probability distribution. Bayes’ theorem is then used to reconcile this prior forecast with a multivariate distribution of observations and their observation errors (a.k.a. a statistical likelihood function) to generate an updated posterior distribution that harmonizes the model and data. Unlike other ensemble approaches, TGEnF and TWEnF include an iterative statistical updating of the process error between the model and the unobserved latent state variables. The TGEnF method differs from the TWEnF in that it uses a normal–gamma conjugacy between the priors of the process error and the likelihood instead of the TWEnF’s multivariate normal–Wishart conjugacy. Because of this, TGEnF doesn’t account for the covariance among model process errors; however, the computational demand is considerably lower, as process errors are estimated independently. This approach also uses a censored normal (a.k.a. tobit) distribution for both the forecast prior and likelihood, which we employ here to ensure carbon and water pools are non-negative. For this analysis, data assimilation was run independently for each site, and so does not consider shared spatial information in the process error covariance or ensemble forecast covariance matrices.

Our overall SDA framework is shown in Figure 3. We first ran 100 SIPNET model (https://github.com/PecanProject/sipnet, Braswell et al., 2005, Supplement 5) ensemble simulations per site, each one year in duration. After the second year, we begin training our ML-Debias model using the forecast residuals (i.e., the forecast minus the observation) and the covariates (Section 2.2.4) from the previous time step. The bias correction is then applied iteratively until 2024, subtracting off the predicted residuals for each variable (Supplement 6). Then, we assimilated the observations into the ensemble forecasts using the TGEnF, the outputs of which became the ICs for the next round of one-year SIPNET forecasts.

**Figure 3.**
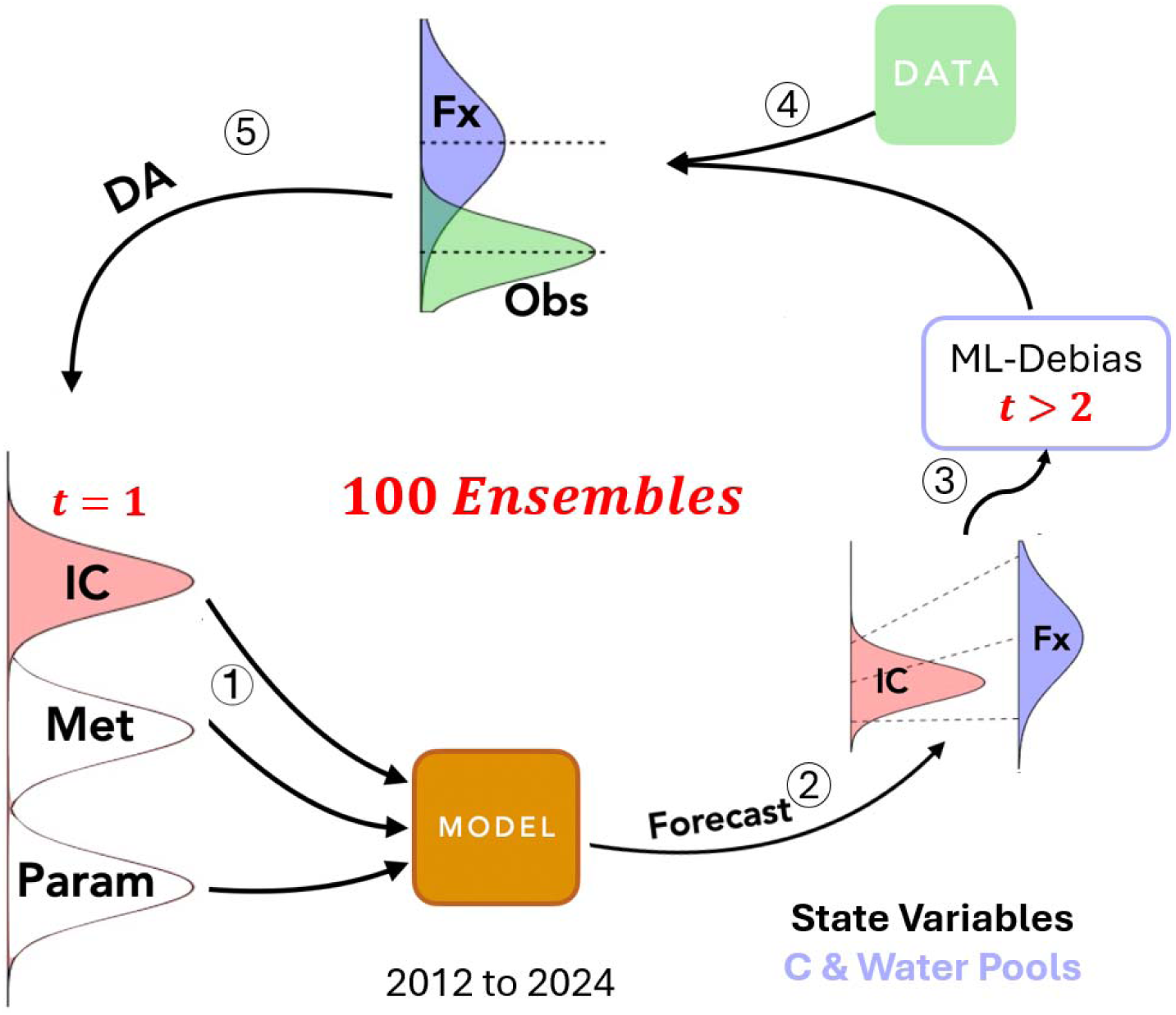
Workflow of the SDA framework. (1) Sample ensemble members from the distribution of data inputs, including initial conditions, ERA5 meteorology drivers, and calibrated parameters. (2) Run an ensemble of the SIPNET model to produce forecasts (Fx) of key ecological states. (3) Apply the ML-Debias algorithm when t is greater than 2. (4) As new data become available for each timestep, (5) assimilate the four data streams (AGB, LAI, soil moisture, and soil carbon) using the TGEnF algorithm, producing harmonized estimates of the carbon and water pools that can be used as ICs for the next model forecast.

### 2.4. Machine-learning emulators

Given the nonlinearity of the SDA system and the need for interpretability, we chose random forest and XGBoost as the machine learning methods to emulate the SDA outputs and interpolate predictions across space. Random forest is a supervised learning method that trains the model by building multiple independent ensemble decision trees. The predictions are obtained by combining the outputs from these decision trees using a bootstrap algorithm (Biau & Scornet, 2016). The XGBoost method, on the other hand, trains the model by building sequential trees, allowing each new tree to correct the errors of the previous one (Chen & Guestrin, 2016). Since the XGBoost approach creates interconnected trees, it may be more prone to overfitting in complex covariate spaces compared to the random forest method. We finally chose Random Forest because it outperformed XGBoost in most cases (Figure S3 in Supplement 7).

We then executed the ensemble ML emulator using covariates in Table 2, as well as the preceding annual average (July 15 to July 14) SMAP SM and MODIS LAI, to capture spatial variability for variables with relatively short temporal memory. Specifically, we fit an ML emulator to each ensemble member, generating 100 maps per time step and variable. Unlike other gridded products (e.g., C estimations derived from remote sensing platforms) that only provide uncertainties for each pixel independently, this approach allows us to estimate covariances across space, time, and pools and to more accurately aggregate uncertainties across these dimensions (e.g., spatial sums).

### 2.5. Analysis and Visualizations

To understand the spatiotemporal patterns of North America’s C and water budgets (Q1), we first calculated maps of ensemble means and SD of each pool across time. To understand the fractions of different carbon budgets, we mapped the ratio of non-photosynthetic vegetation (stem + root) to the entire carbon stock (leaf + stem + root + soil carbon). We then evaluated the contributions of different covariates to C pool patterns by plotting the variable importance within the ML emulator averaged across time and ensemble. To assess temporal trends in the mean and uncertainty of each variable over time (from 2015 to 2024, to avoid impacts of ICs) we fit pixel-level linear regressions, removing slopes with p-values < 0.05, and then mapped the slopes across NA. The uncertainty for each variable across 8,000 locations were compared between SDA outputs and the corresponding datasets used in this study.

To assess the spatiotemporal accuracy of our C estimations (Q2) and the temporal improvements from the ML debiasing method (Q3), for the 8,000 selected sites, we first calculated R^2^and RMSE values over time between our SDA analysis and the corresponding data constraints. Afterwards, we plotted the residual errors between our emulated maps and the data constraints.

In addition to comparing the SDA output to the data constraints, we also validated it against held-out observations for AGB (GEDI, Dubayah et al., 2022; ICESat-2, Duncanson et al., 2023; and USFS FIA BIGMAP, Wilson et al., 2018), where GEDI and ICESat-2 are LiDAR (Light Detection and Ranging) platforms, and BIGMAP is based on Landsat pixels and FIA inventory data from 2014 and 2018. During the comparison, we first scaled the ICESat-2 AGB from 30 m to 1 km and then combined the GEDI and ICESat-2 AGB maps. We then compared AGB across 8,000 sites, first analyzing the entire NA, then dividing into different latitude ranges (>50°, 30°-50°, and 7°-30°) and forest types (deciduous and evergreen forests). After that, we aggregated both the SDA analysis and emulated maps from 2014 to 2018 to make fair comparisons after upscaling the FIA BIGMAP AGB to 1-km resolution.

We also validated our SOC estimates against the ISCN records. Although the ISCN database is partially used in the SoilGrids products (Poggio et al., 2021), it remains valuable for assessing the accuracy of our estimates against actual soil core measurements. First, we filtered the ISCN observations by soil depth and aggregated them by EPA level 2 ecoregion classifications. Then, we calculated the mean SOC estimates per ecoregion and compared them with our SDA analysis using the same aggregation approach. Confidence intervals were obtained by quantifying the 0.025 and 0.975 quantiles of the SDA ensembles across ecoregions.

Finally, to show the improvements in accuracy in C estimates from the ML debiasing method (Q3), we also conducted continental SDA experiments without the ML debiasing algorithm and compared AGB and SOC estimates against the same set of data products as above (Supplement 8, Figures S4 - S7). After that, we evaluated the contributions of different covariates to model biases for each pool by plotting the average variable importance of the ML debiasing routine averaging over time and ensemble.

**Table 5.**
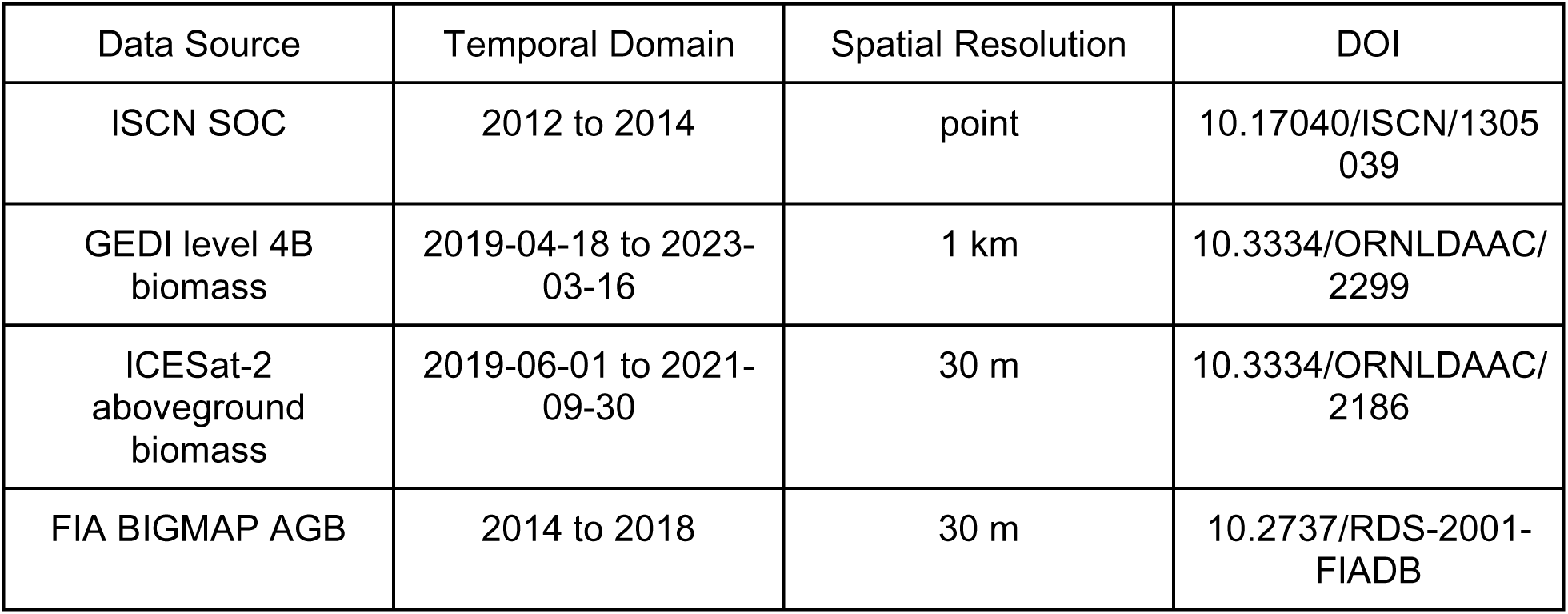
Data used for validation, summarized by name, temporal domain, spatial resolution, and DOI.

## 3. Results

### 3.1. Carbon and water budgets

#### 3.1.1. Mean and uncertainty at 1km resolution

The July 2024 AGB and LAI maps exhibit similar spatial patterns, with the highest values in forested regions such as the Pacific Northwest, the East Coast, and Central America (Figure 4). LAI is also high throughout the agricultural regions of the Midwest US and the Canadian Prairie Provinces, and lowest in drylands and tundra. SM exhibits a similar spatial pattern to aboveground biomass, with higher biomass density generally corresponding to higher SM. The high peak in the SM distribution indicates tropical and arctic peatlands with saturated organic soils. The SOC shows an increasing trend from south to north across NA. The SD distributions exhibit a similar spatial pattern to the mean map of each variable (Figure 5), indicating that for all carbon pools, uncertainties were proportional to pool sizes (Figure S8): LAI (0.147, p-value < 0.001), SOC (0.119, p-value < 0.001), and AGB (0.072, p-value < 0.001). The non-photosynthetic plant C map (Figure 6) shows that wood carbon dominates the C budget of the major forested ecoregions, particularly western rainforests and eastern forests, while SOC dominates the entire carbon stock across the Northern Arctic, Great Plains, and Southern Deserts.

**Figure 4.**
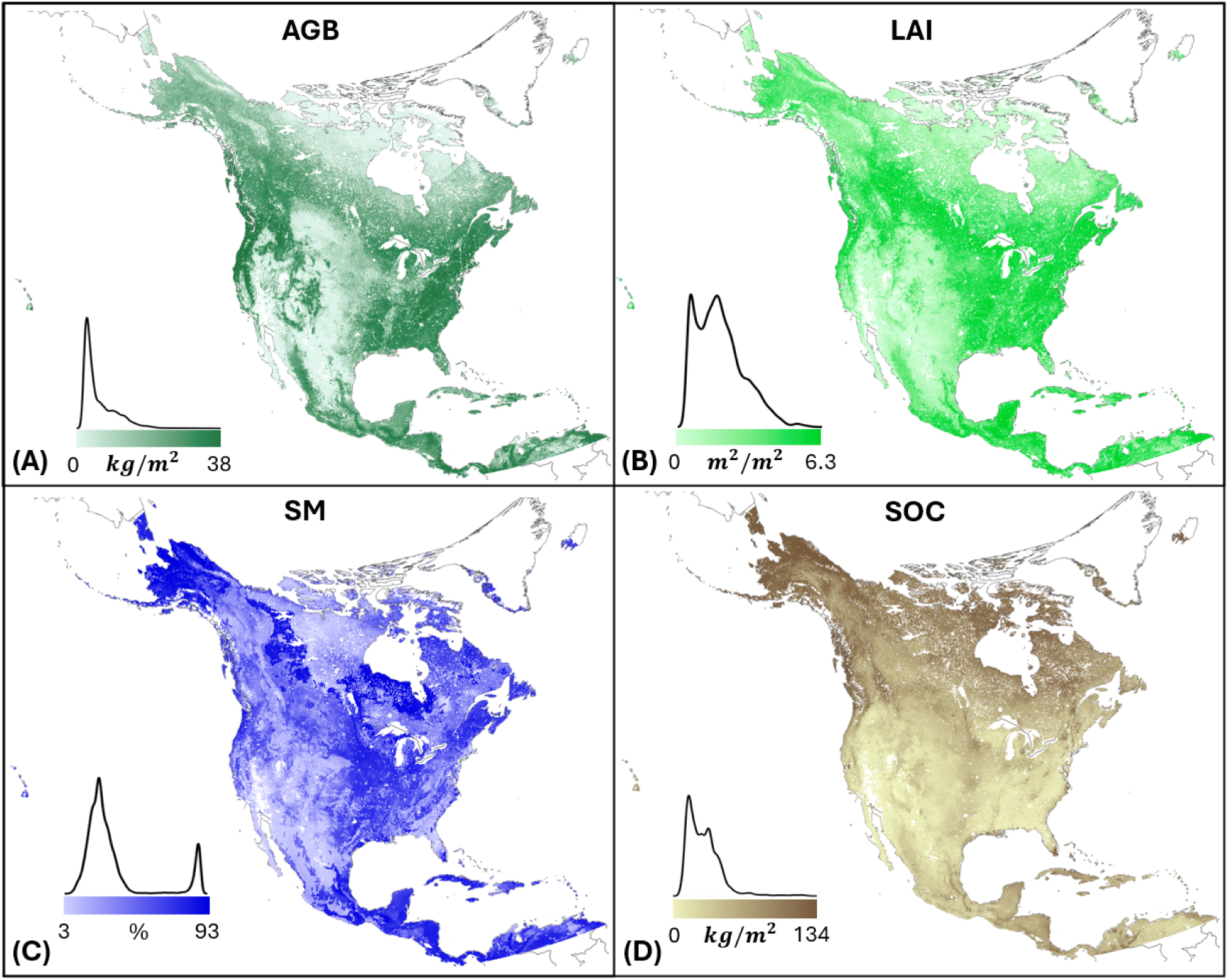
The spatial patterns of the ensemble means (n=100) for (A) AGB, (B) LAI, (C) SM, and (D) SOC for July 15, 2024. The corresponding density distributions are plotted for each variable with the color bar and range listed below.

**Figure 5.**
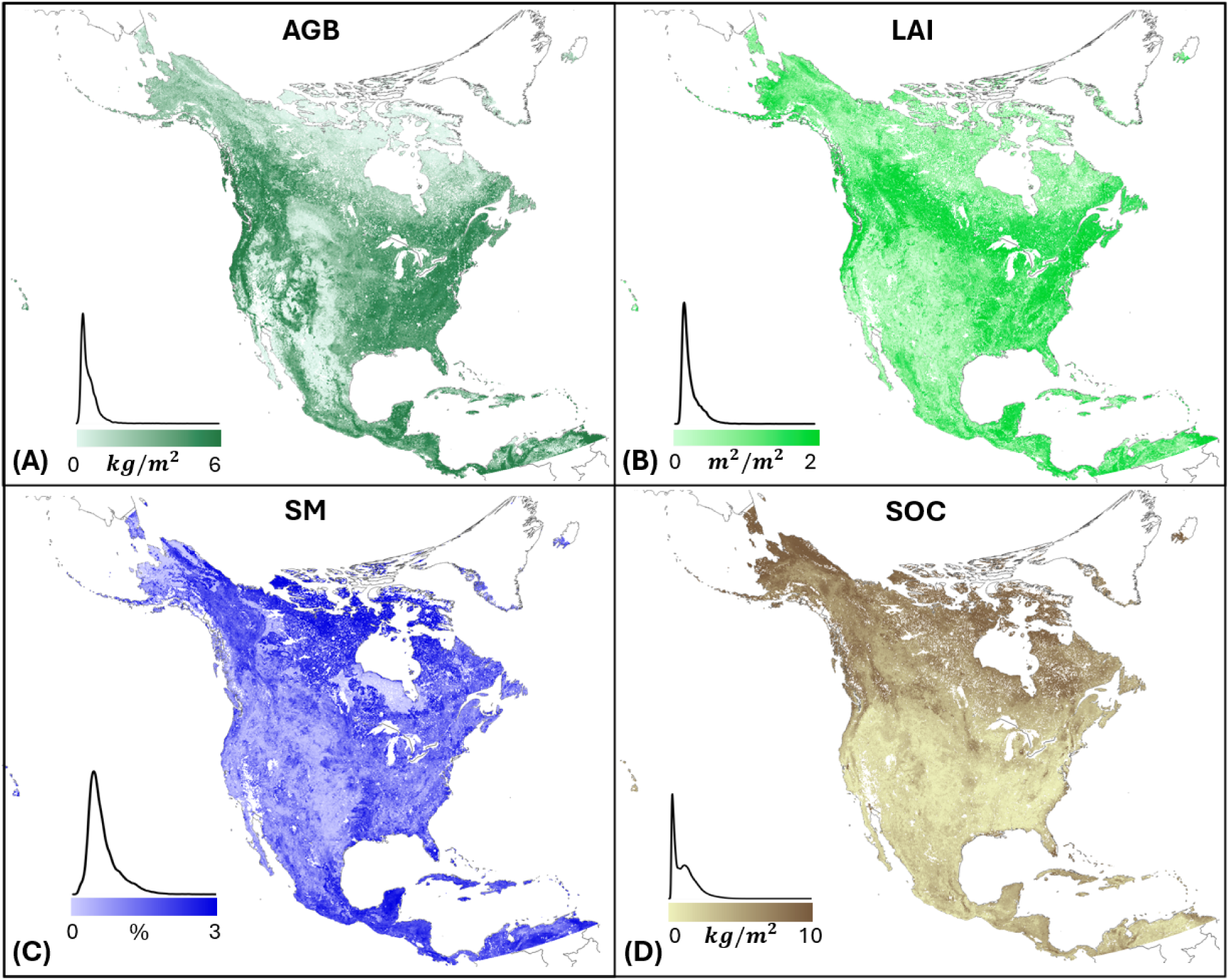
The spatial patterns of the ensemble standard deviation for (A) AGB, (B) LAI, (C) SM, and (D) SOC. The corresponding density distributions are plotted for each variable with the color bar and range listed below.

**Figure 6.**
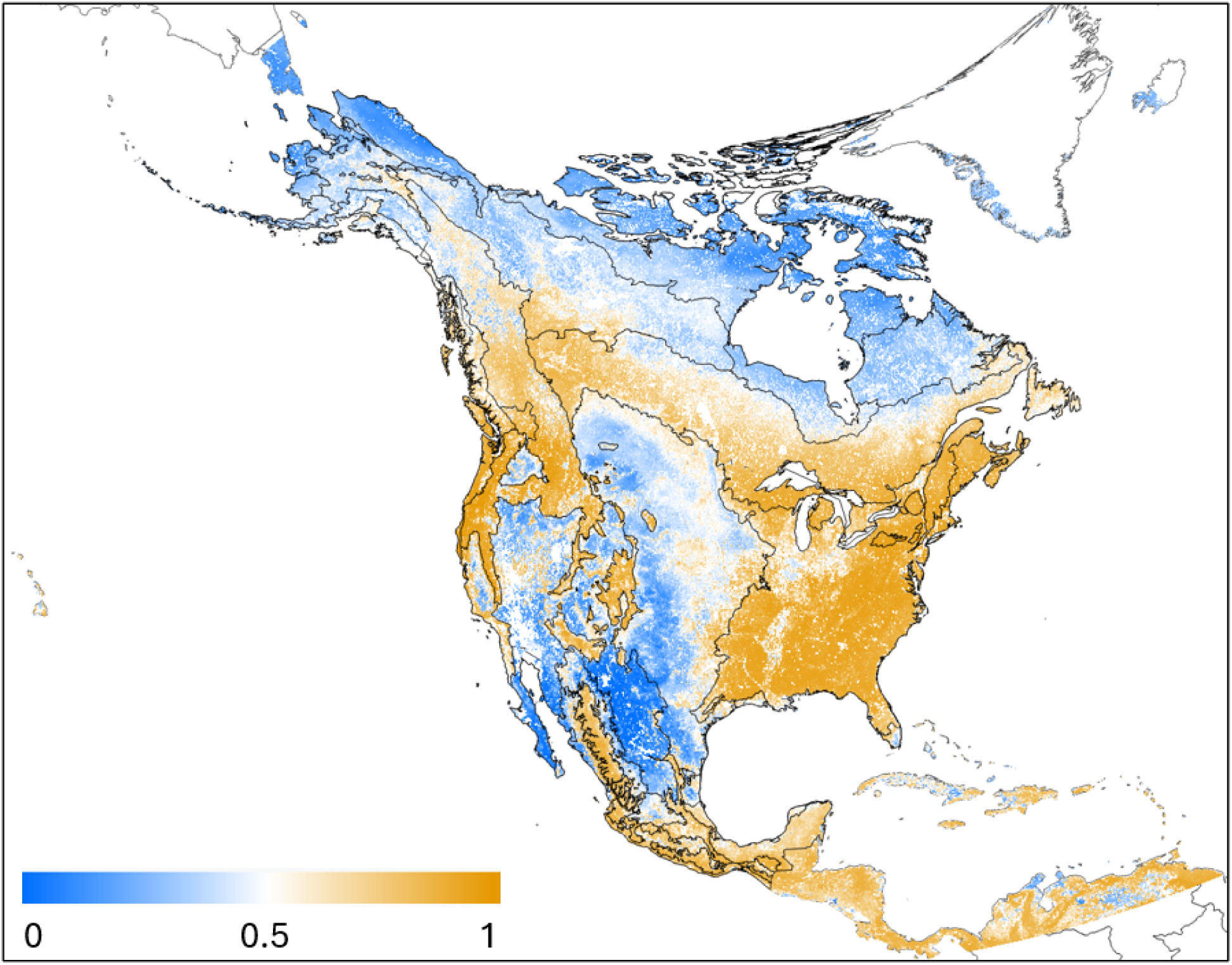
Proportion of non-photosynthetic plant C to the entire carbon stock. Here, we calculated the ratio of (stem + root) to the entire carbon stock (leaf + stem + root + soil carbon) for the 2024 predictions.

The ML emulator variable importance analysis suggests a consistently high importance of the corresponding covariate within each pool (AGB to AGB pool, LAI to LAI pool, etc., Figure 7). In contrast, the remaining covariates show similar but slightly different patterns of importance. For example, land cover and temperature are important in AGB predictions; AGB, land cover, and climate variables (solar radiation, temperature, and vapor pressure) are important in LAI predictions; precipitation and percent sand are important for SM predictions; and elevation, soil pH, precipitation, and solar radiation are important in SOC predictions.

**Figure 7.**
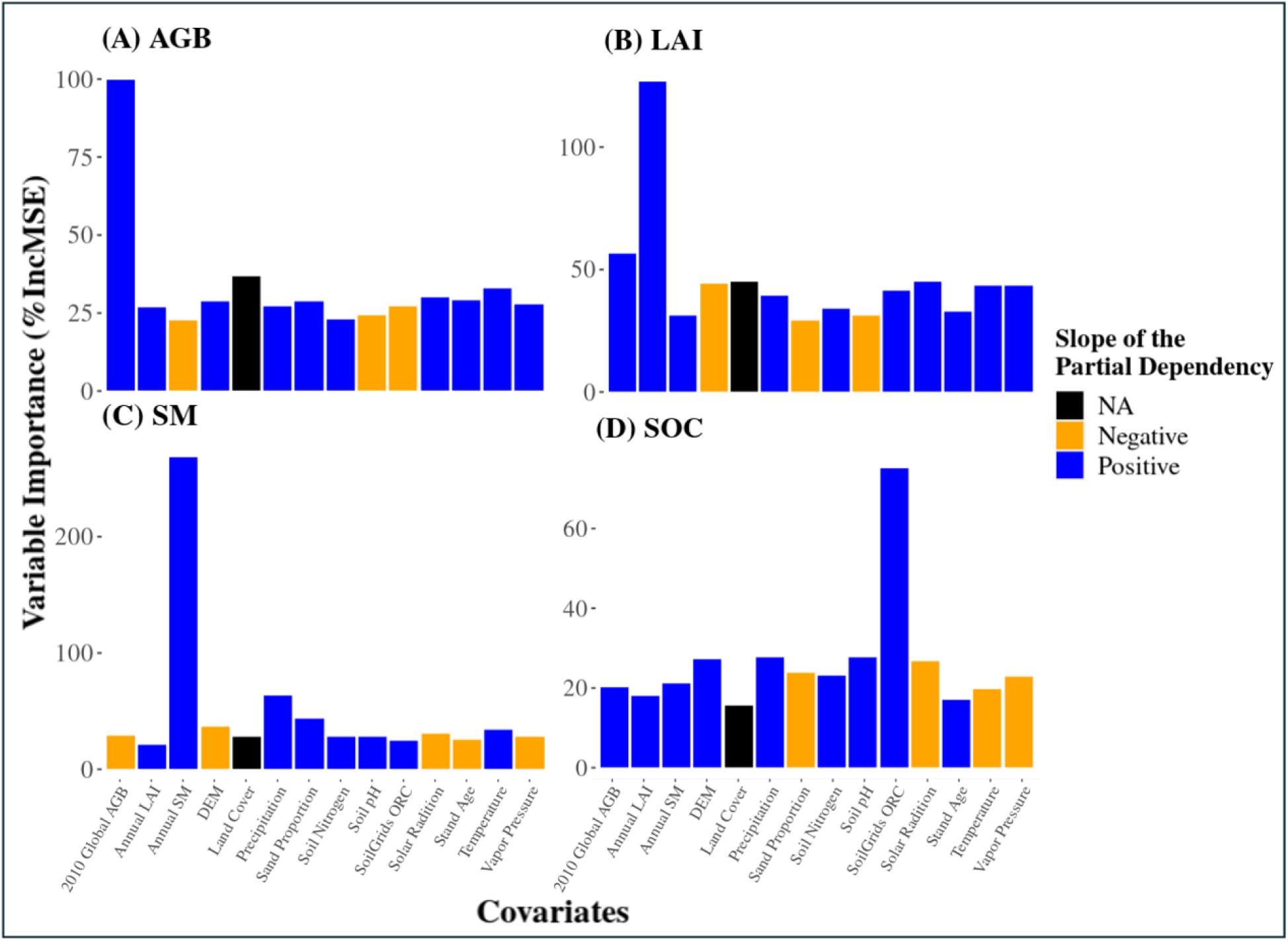
The variable importance (percentage increase in Mean Squared Error, %IncMSE) of covariates for (A) AGB, (B) LAI, (C) SM, and (D) SOC during the ML emulator in 2024. The colors represent the slope of the partial dependency trends (Landcover is assigned NA because it’s not continuous variable).

The temporal trends of each variable (Figure 8) indicate that we generally predict a slight increase in AGB over time in the Alaskan tundra, decreases across the western US forests, and mixed responses across the eastern CONUS. LAI shows an increasing trend across the tundra and agricultural Great Plains ecoregions, while decreasing across the Canadian boreal and west-coast US forests. SM decreases over time across eastern Canada and parts of the eastern US, while increasing across tropical Central America and the Caribbean. SOC remains stable across most regions, except for a slight increase in Alaska. The uncertainty trends (Figure 9) indicate that the AGB uncertainty generally increases for most forested ecoregions, with relatively higher magnitude in eastern forests. By contrast, LAI uncertainty generally declines, with the largest decreases in boreal and western forests that experienced LAI declines, and sporadic increases across the central US. The SM uncertainty change is also relatively small but slightly increases in patches over time across the northern Great Plains and Canada. Finally, the SOC uncertainty decreases throughout the entire period, with greater reductions at higher latitudes.

**Figure 8.**
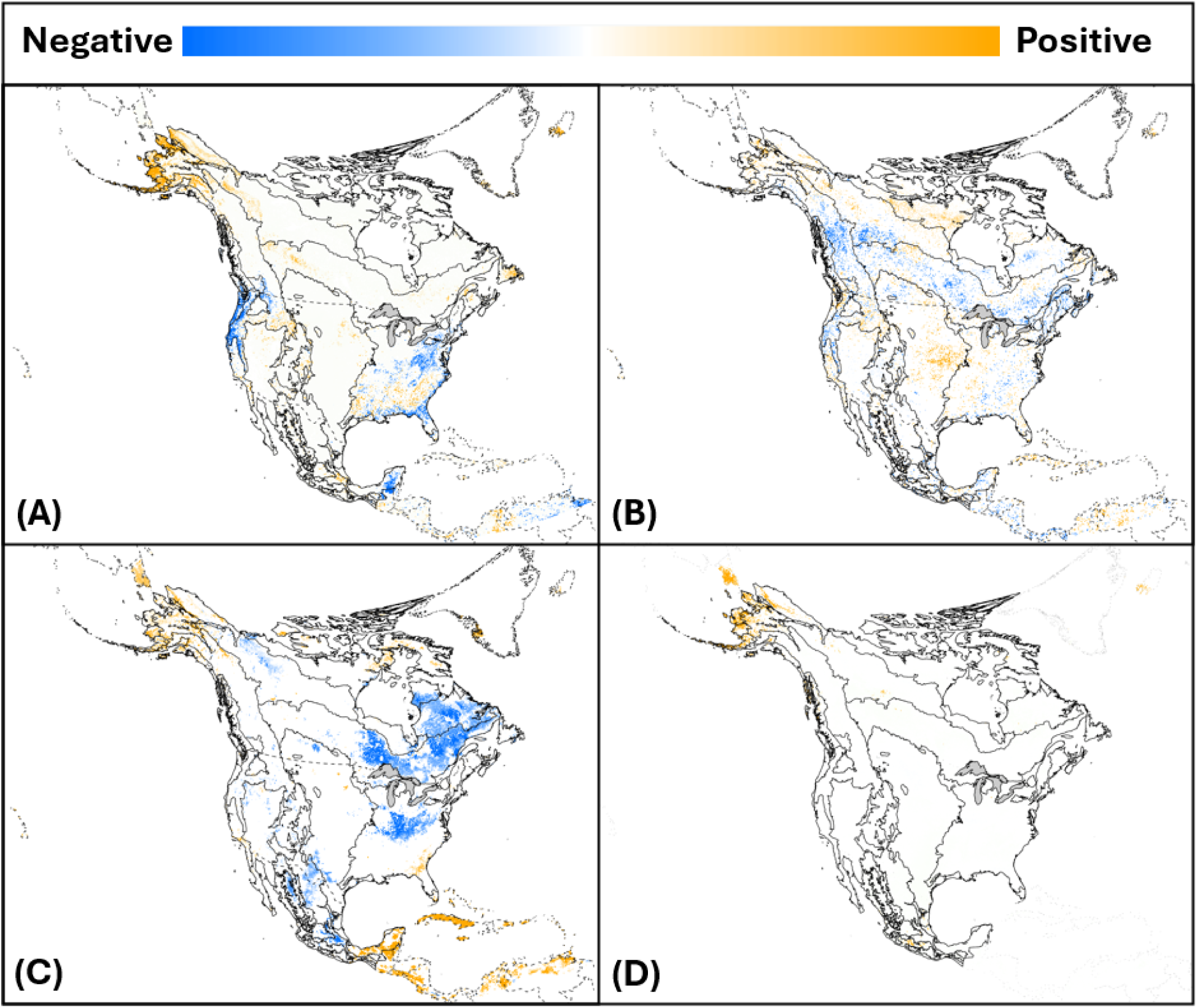
Pixel-level regression slopes for change across time (2015-2024) for (A) AGB (from -1 to 0.3), (B) LAI (from -0.4 to 0.34 LAI/year), (C) SM (from -5 to 5 % of volumetric soil water), and (D) SOC (from -1 to 1). Pixels with p-values < 0.05 were removed. Background polygons represent Level-1 EPA ecoregions.

**Figure 9.**
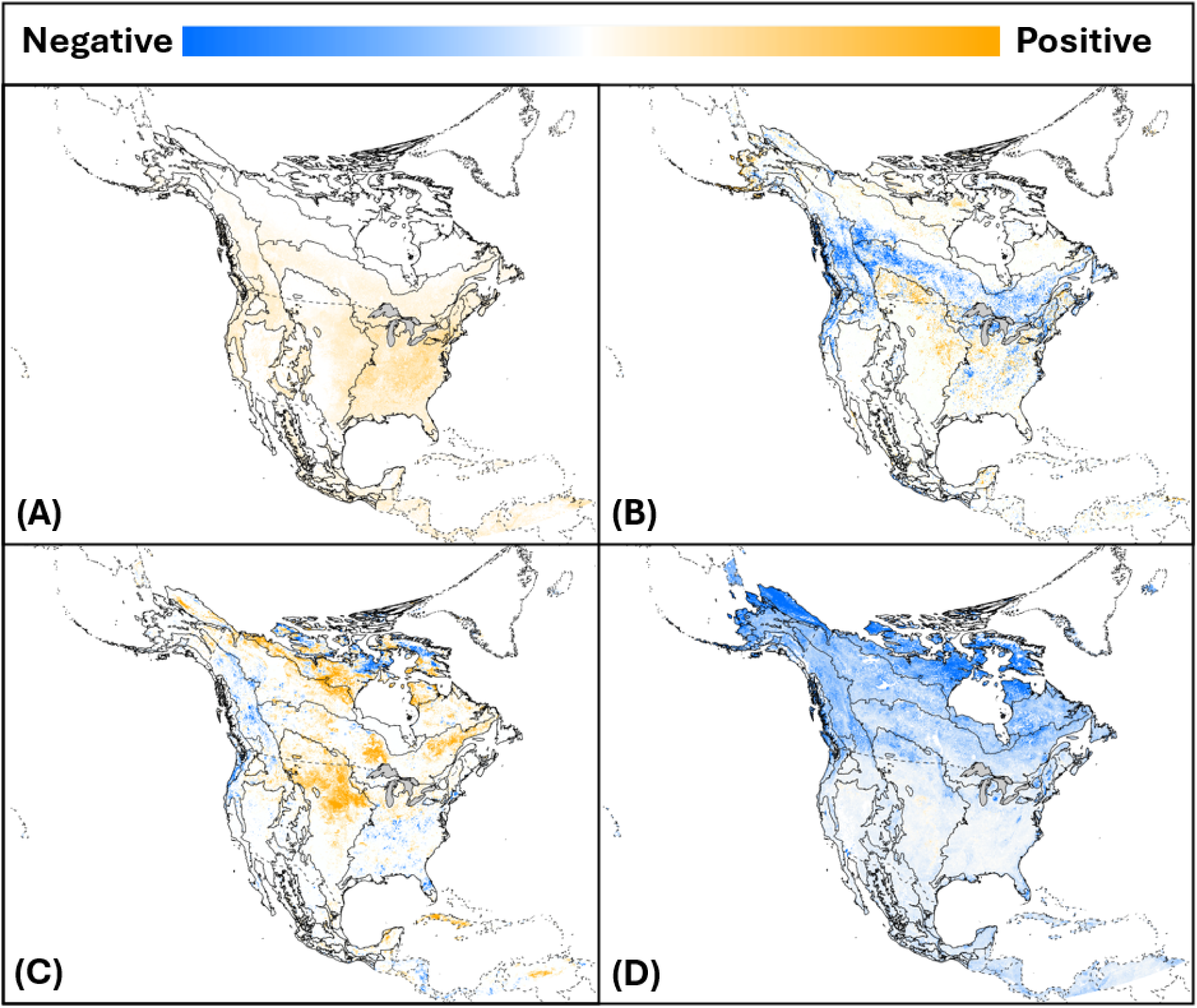
Pixel-level regression slopes for change in uncertainty across time (2015-2024) (A) AGB (from -0.12 to 0.22), (B) LAI (from -0.1 to 0.1 LAI/year), (C) SM (from -0.2 to 0.2 % of volumetric soil water), and (D) SOC (from -1.7 to 0.1). Pixels with p-values < 0.05 were removed. Polygons represent Level-1 EPA ecoregions.

### 3.2. Comparisons against Data Constraints

Using the SDA + ML-debias algorithm, we observed a decrease in error metrics from 2012 to 2024 for both AGB (RMSE from 1.28 to 0.79, reduced by 38.3%) and SOC (RMSE from 8.575 to 1.80, reduced by 79%) (Figure 10), then remained stable afterwards, though with slight reductions in AGB R^2^ and increased RMSE around 2018, and a slight downward trend in SOC RMSE from 2016 to 2024. By comparison, LAI and SM showed a consistent high R^2^ and low RMSE across the entire time period.

**Figure 10.**
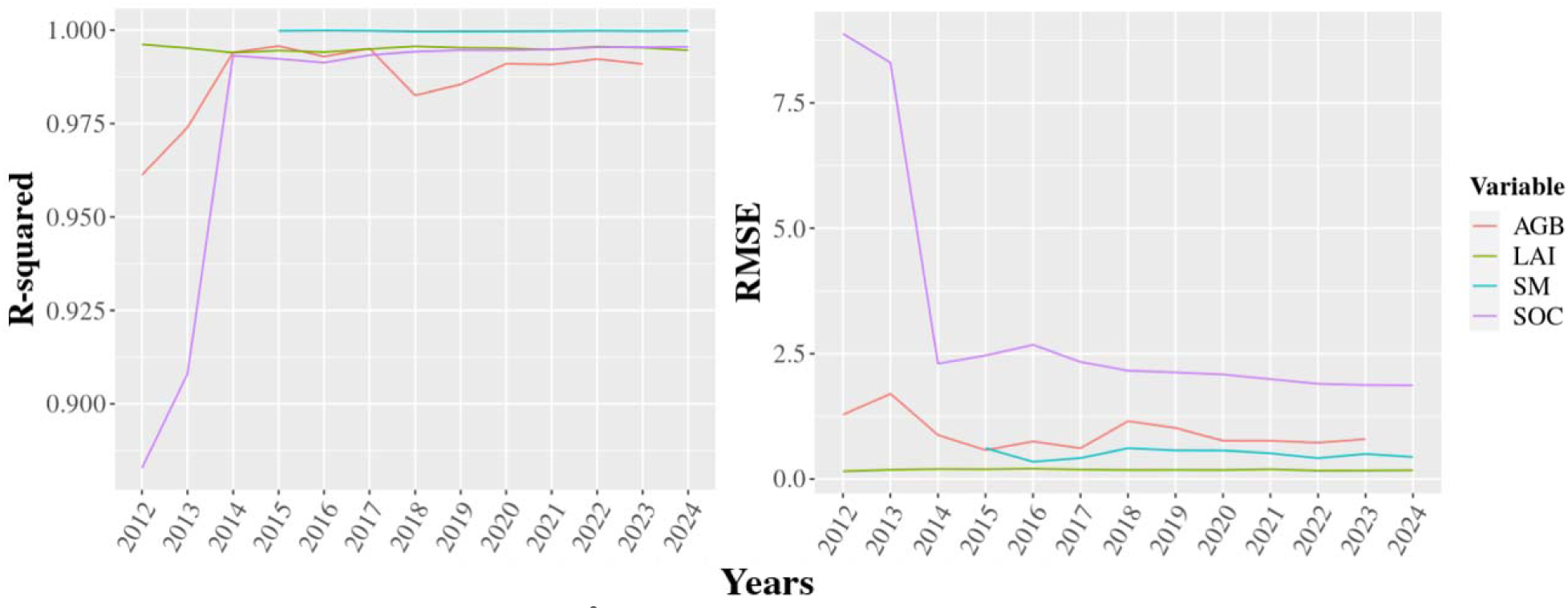
The temporal trends of R^2^ and RMSE from 2012 to 2024 across C and water budgets for the pre-selected 8,000 locations. Any missing values represent NAs in the observation (e.g., SMAP SM is not available until 2015).

Although AGB shows high accuracy relative to observations, the largest residual errors occurred along the western US forested coast (Figure 11). LAI and SM residual maps indicate roughly random distributions, with a slight LAI overestimate across Alaska and an underestimate for western temperate rainforests. The SOC residuals (-4 to 4) show smaller variations than AGB (-10 to 10) with the higher errors distributed in higher latitudes, particularly in western Canada and Alaska (Figure 11).

**Figure 11.**
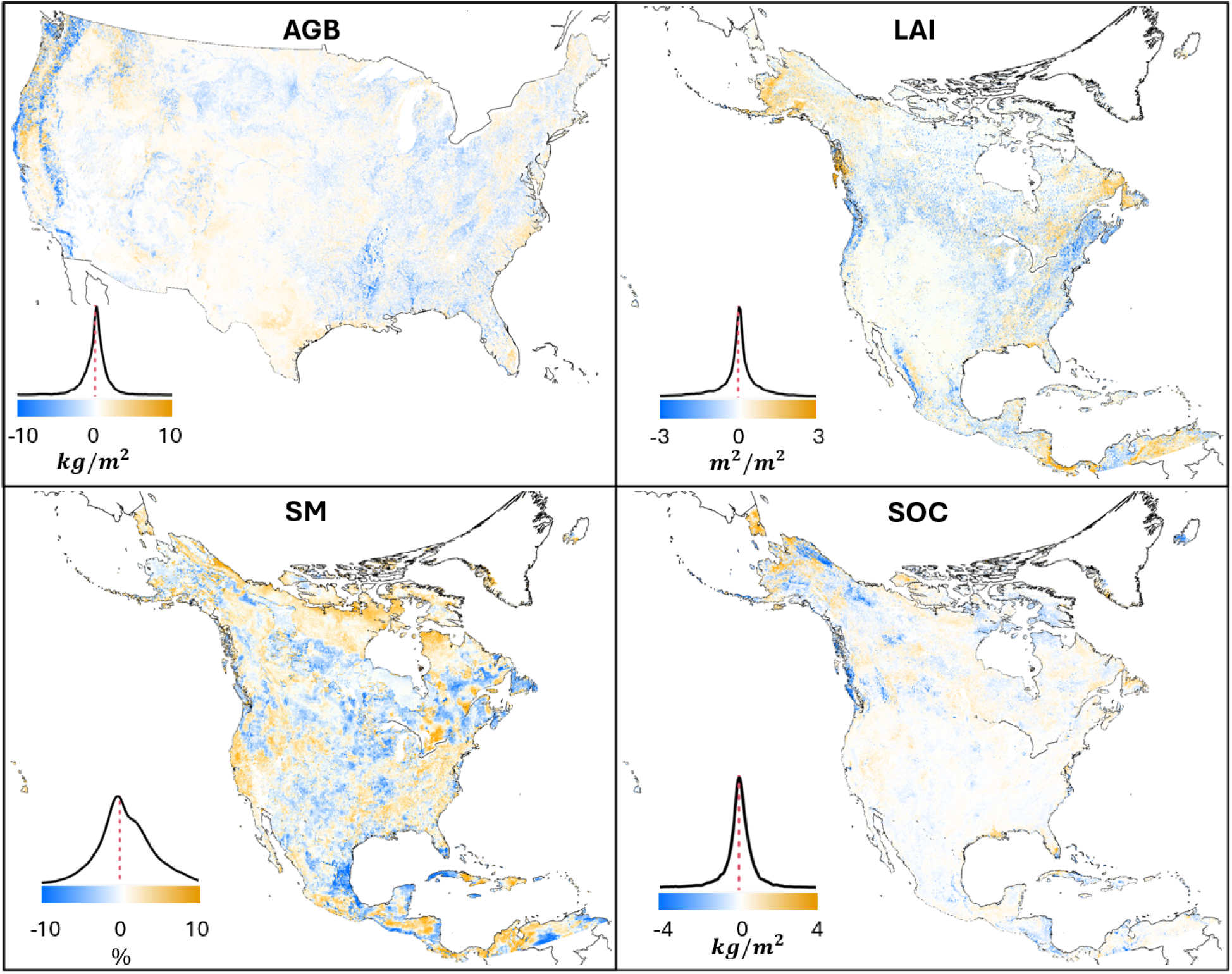
Spatial patterns of the residual differences between the ensemble mean of the SDA results and the corresponding data constraints for (A) AGB, (B) LAI, (C) SM, and (D) SOC. The corresponding density distributions are plotted for each variable with the color bar and range listed below.

SDA benefits from both comprehensive uncertainty propagation and the reduction of uncertainties due to borrowing strength across a process model and multiple data constraints. We assess the magnitude of this effect by comparing the distribution of pixel-level uncertainties between our SDA analysis and the assimilated data products. Figure 12 shows that the AGB uncertainty has been reduced by 82.4% compared to the assimilated LandTrendr product, displaying a similar pattern to the held-out GEDI and ICESat-2 AGB products. The uncertainty of LAI similarly decreases by 37.5% compared to the MODIS LAI products. The soil water uncertainty is roughly the same between the analysis and SMAP. Finally, SOC uncertainty has also been substantially reduced by an average of 77.0% relative to the SoilGrids estimates.

**Figure 12.**
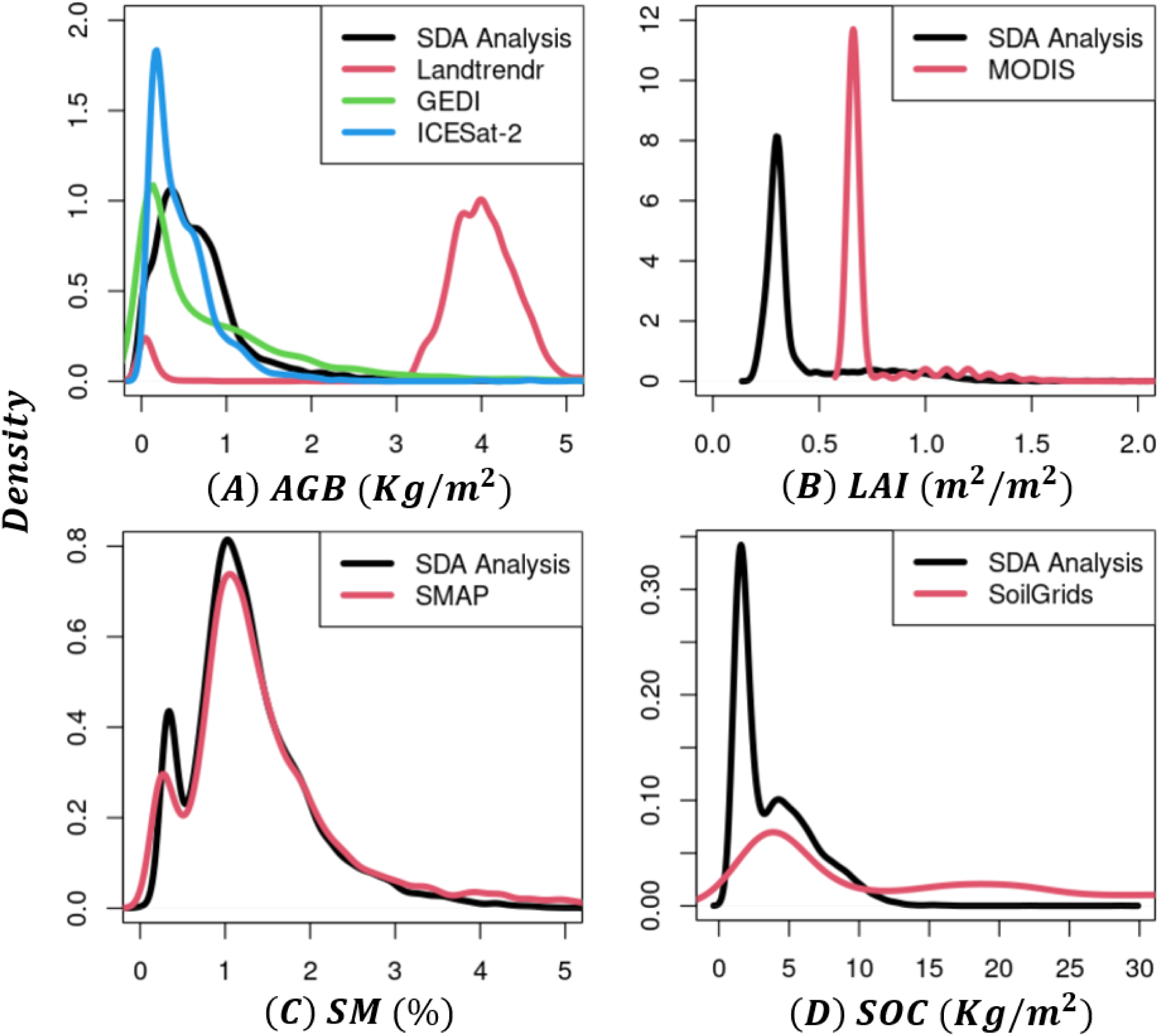
Uncertainty comparisons for the 8,000 sites between our SDA outputs and the data constraints used in the SDA workflow for (A) AGB, (B) LAI, (C) SM, and (D) SOC.

### 3.3 AGB Validation

In addition to comparing against the data constraints used in the SDA workflow, it is also essential to evaluate the SDA against independent held-out data products. Here, we first validate our results against the ICESat-2 + GEDI AGB products. Comparing the averaged AGB analysis from 2019 to 2023 at 8,000 pre-selected locations against the extracted AGB estimations from the ICESat-2 + GEDI AGB, we obtain an R^2^ of 0.73 and RMSE of 3.51 kgC/m^2^ (Figure S9), with most points spreading along the x = y line, with denser regions of points indicating areas with more agreement.

Dividing the data by latitude and forest type reveals that the SDA generally achieves high accuracy, with a high density of points along the one-to-one line; however, forests with an AGB greater than ∼15 kg/m² tend to be underestimated compared to the ICESat-2 + GEDI estimates (Figure 13). In general, the SDA was more accurate in evergreen forests than in deciduous forests, particularly at lower latitudes. The exception to this pattern is high-latitude forests, where deciduous and evergreen forests perform similarly. This region tends to have lower biomass, which is the condition where the SDA tends to be more accurate. In regions where the LandTrendr product is assimilated (roughly between 50° and 30° N), the SDA AGB underestimations are consistent with underestimates in the LandTrendr product itself, which may experience saturation problems over high-biomass forests (Figure S10). These underestimations are more pronounced in high-biomass forests below 30° N, with deciduous (respectively evergreen) forests underestimated when AGB exceeds approximately 5 kgC/m^2^ (resp. 10 kgC/m^2^). Maps of residual error (Figure 14) display similar patterns, indicating higher accuracy above 50°N. However, the SDA underestimates AGB in eastern deciduous forests and western temperate rainforests between 50° and 30°N, as well as tropical and subtropical evergreen forests below 30°N.

**Figure 13.**
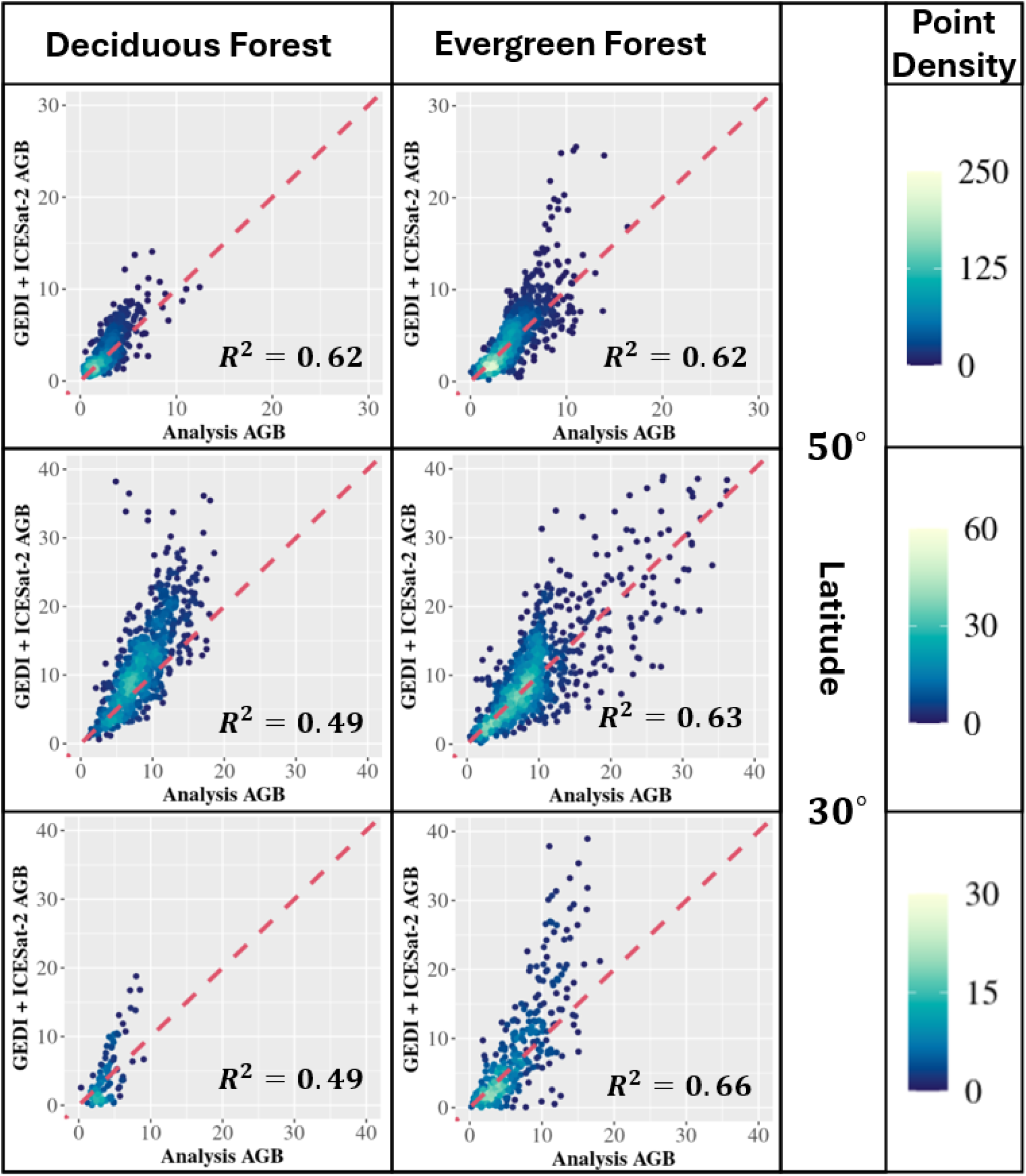
Comparisons between GEDI + ICESat-2 and SDA AGB split by latitude range (> 50 degrees, 30 - 50 degrees, < 30 degrees) and forest types (deciduous and evergreen forest, respectively), where the 1:1 lines are labeled as red dashed lines. The point density colors the dots: yellower means more points are clustered, while bluer means fewer points are clustered.

**Figure 14.**
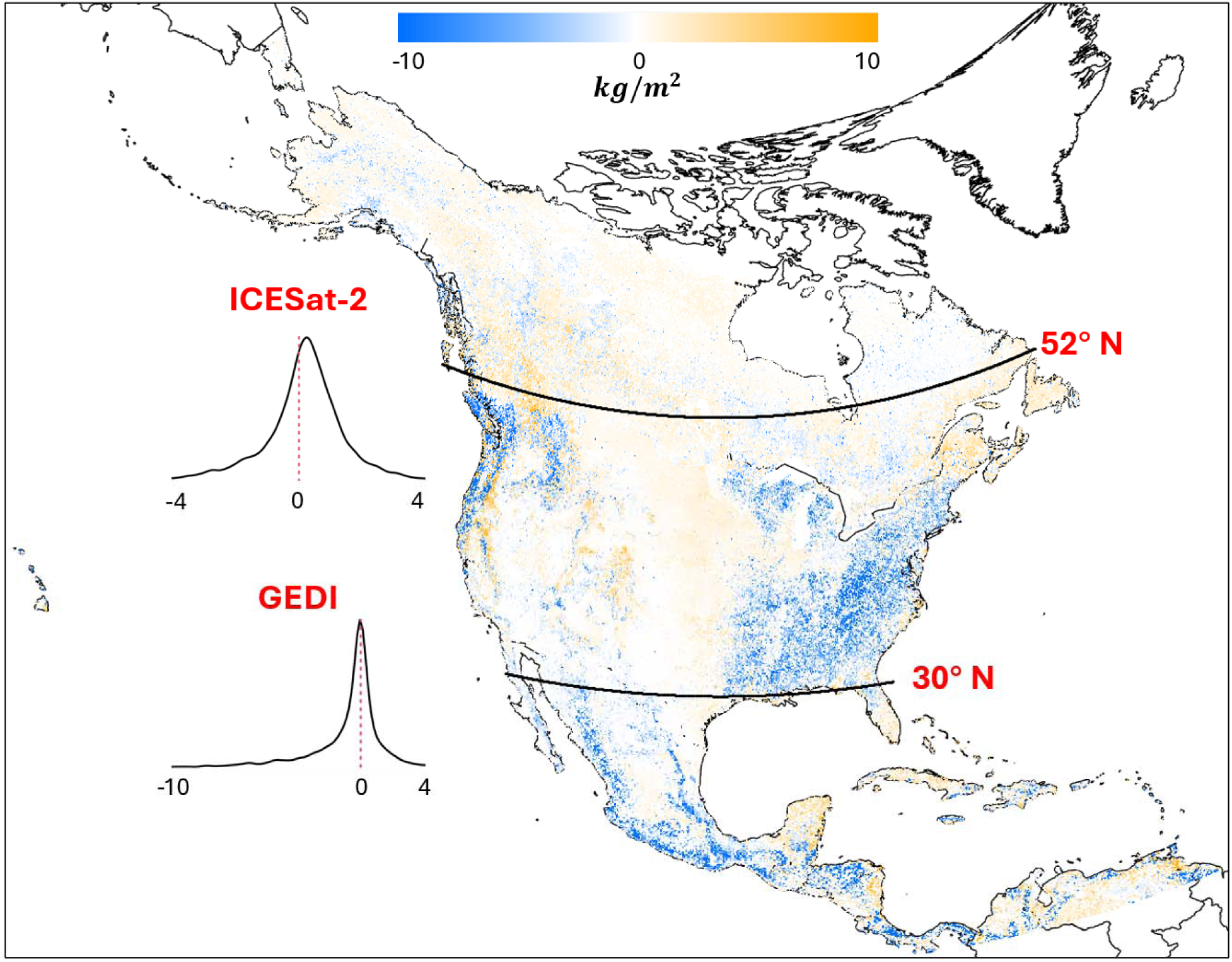
The spatial patterns of the differences between the SDA AGB analysis mean and the ICESat-2 and GEDI products. The corresponding density plots show the distributions of residual errors for ICESat-2 and GEDI AGB products. The 2.5%, 50%, and 97.5% quantiles are, respectively, -3.06, 0.24, and 2.77 for ICESat-2, and -10.4, 0.18, and 3.97 for GEDI.

We also validated the SDA results, averaged over 2014-2018, against the FIA BIGMAP database, achieving high accuracy ( = 0.74; Figure 15). Similar to the ICESat-2 + GEDI validation, we found that AGB accuracy in deciduous forests (R^2^ = 0.54, with a slight underestimation of 0.69, p-value < 0.001, relative to the FIA estimates) is lower than in evergreen forests ( = 0.8, p-value = 0.47). The spatial map of residual error shows a pattern similar to that of the point-wise comparisons (Figure S11).

**Figure 15.**
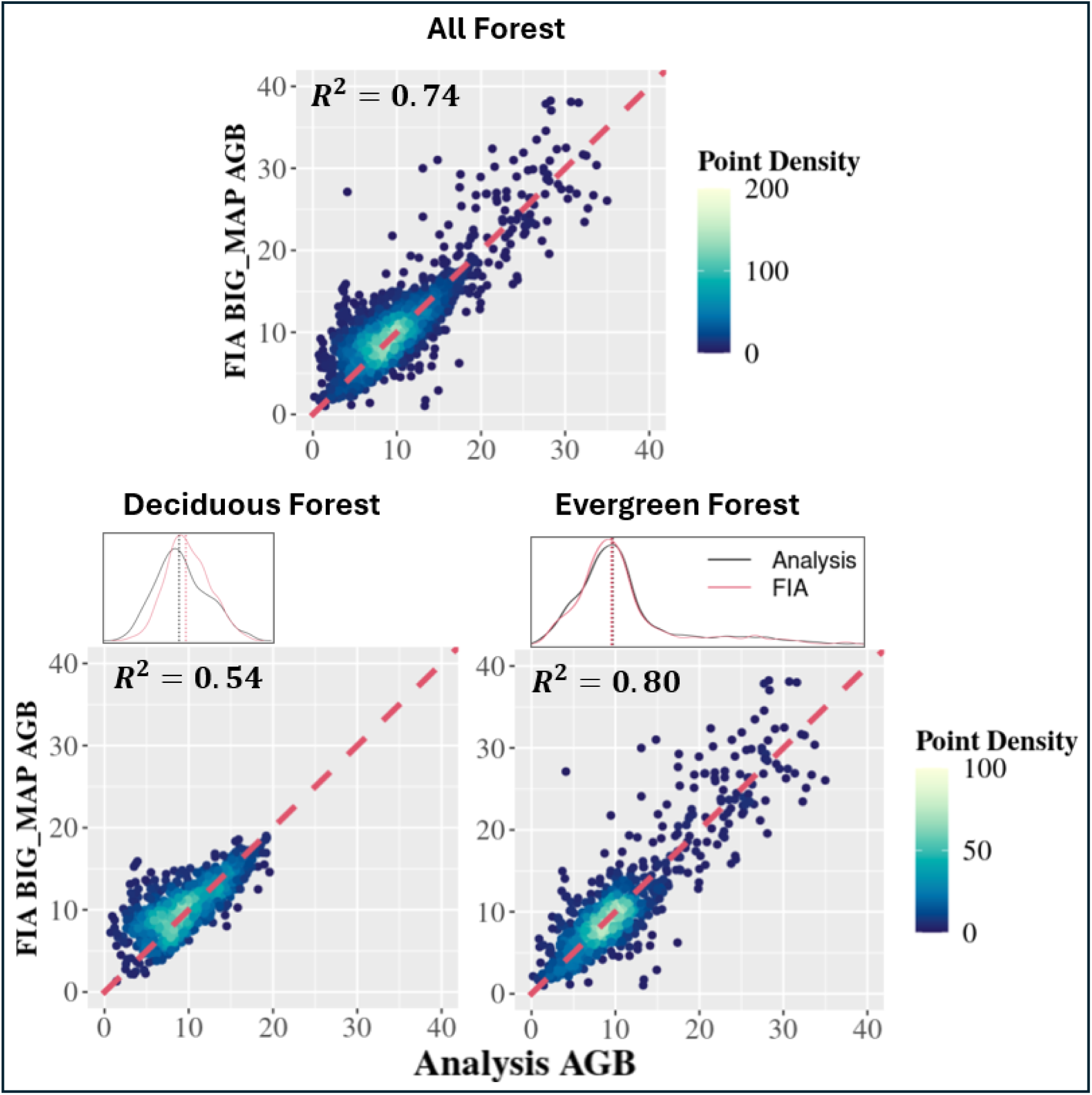
Comparisons between the SDA AGB analysis (averaged from 2014 to 2018) and the FIA BIGMAP database across (A) all forest, (B) deciduous forest, and (C) evergreen forest.

### 3.4 SOC validation

Due to the spatial misalignment between the 8,000 SDA SOC estimates and ISCN records, both datasets were aggregated to EPA Level 2 ecoregions for validation (Table S1). Most SOC estimates align well with the ISCN measurements and fall within the upper and lower bounds, with an R² of 0.44, except in ecoregions with limited SOC data, which are indicated by dark points outside the green boundaries (Figure 16). The outliers occur within the following ecoregions: Taiga shield, with one observation; Boreal plain, with two observations; and Tamaulipas-Texas semiarid plain, with 65 measurements.

**Figure 16.**
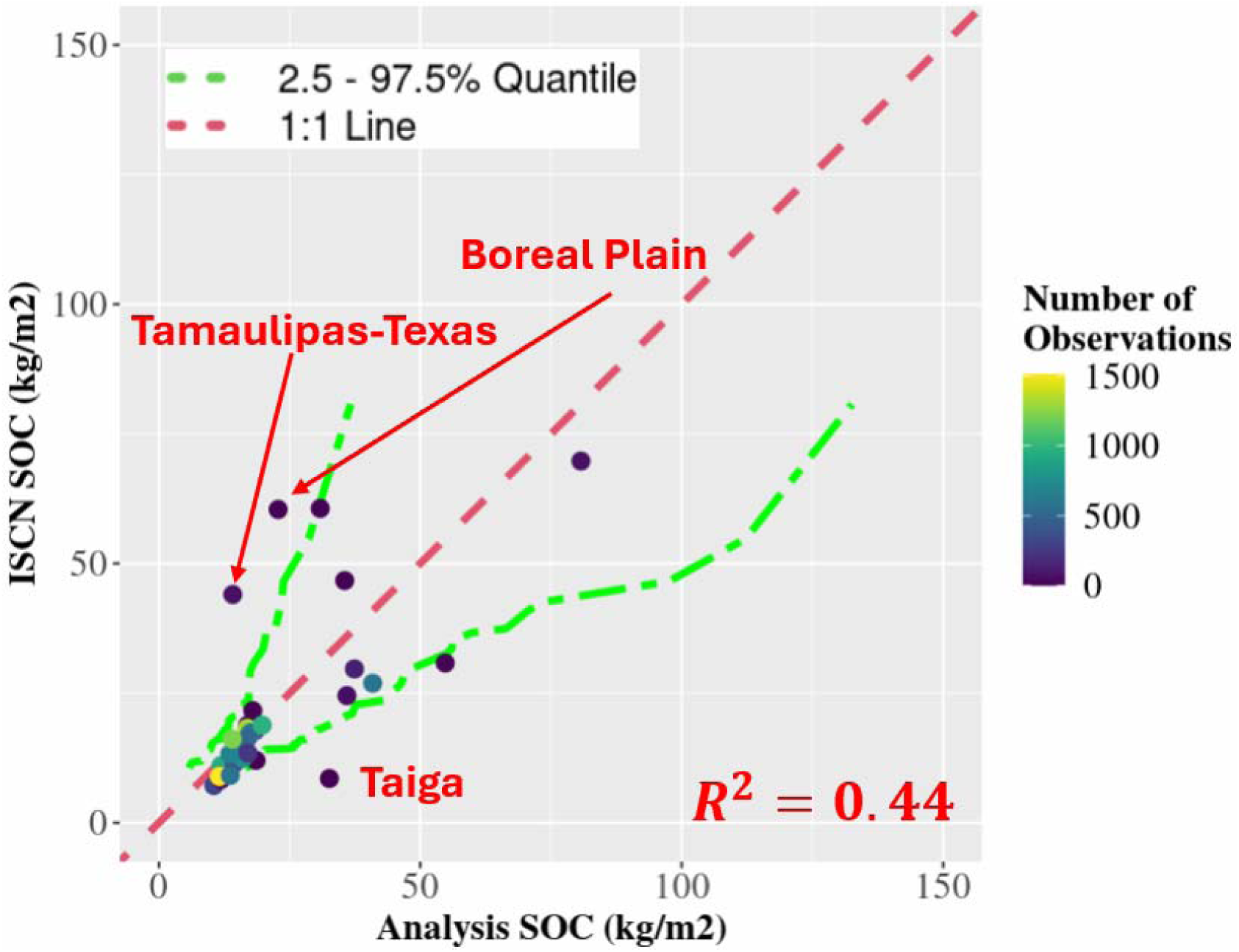
Comparisons between ISCN and the SDA SOC Analysis across the level 2 EPA ecoregions, with quantiles (2.5, 50, and 97.5%) from the ensemble analysis results. Points are colored by the number of available ISCN records for each ecoregion.

### 3.5 ML debiasing

To compare the major C budgets before and after applying the ML debiasing, we regenerated results from the previous sections without debiasing. From these results, we observe that ML debiasing led to a substantial improvement in SOC, with a 50% reduction in the RMSE from 4.9 to 1.8, an even larger RMSE reduction of 66% in AGB (from 1.63 to 0.79), but negligible improvements in the already highly accurate LAI and SM (Figure S4). ML debiasing also led to improved accuracy compared with the FIA BIGMAP database, with the R² value increasing from 0.71 to 0.74 (Figures S6 and 15), primarily due to improvements in evergreen forests. Regarding the GEDI + ICESat-2 validations, the overall R-squared values remain the same (0.73), but with improved accuracy across the northern (from 0.56 to 0.62 and from 0.58 to 0.62 for the deciduous and evergreen forests, respectively) and southern evergreen forests (from 0.58 to 0.66) (Figures S5 and 13). The ISCN validation improved with the debiasing algorithm, as evidenced by the reduction from six to three outliers (Figures S7 and 16).

Residual errors are primarily explained by that year’s model forecast (Figure 17). This was followed by the previous year’s lagged residual and previous observations of the focal variable being either the second or third most important predictors (e.g., lagged error was second for the more strongly corrected AGB and SOC, while prior observations were second for the lower error LAI and SM). For LAI error there were also moderate contributions from the AGB initial conditions, and land cover.

**Figure 17.**
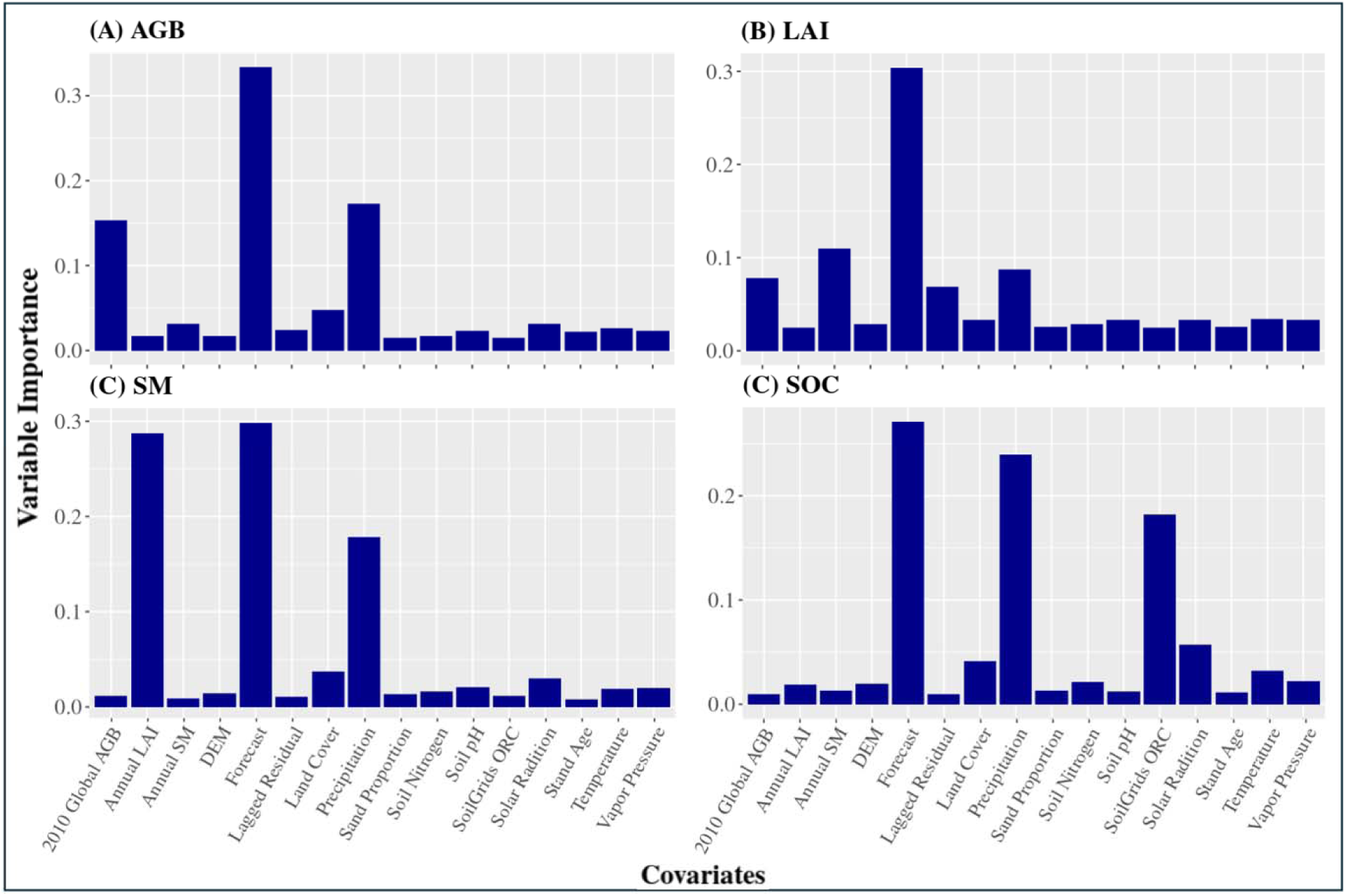
The variable importance (proportion of variability explained by each covariate) of covariates for (A) AGB, (B) LAI, (C) Soil Moisture, and (D) SOC during the ML-debias in 2024.

## 4. Discussion

In this study, we applied our novel hybrid SDA–ML framework to estimate key C budgets and the associated uncertainties across North America from 2012 to 2024. We demonstrated the feasibility of a continental-scale reanalysis for carbon MRV at a moderate spatial resolution (1km) and with reduced uncertainty relative to the corresponding data constraints, particularly for the larger magnitude and higher-memory AGB and SOC stocks where uncertainties were reduced by 83% and 77%, respectively. This reanalysis will be used to answer three key research questions as follows.

### 4.1. What are the general spatial-temporal patterns and trends of C budgets and the associated uncertainties, and their spatial-temporal correlations to explanatory covariates?

Overall, the four key C and water pools are well predicted and align with established biogeographic patterns (Figure 6), while the uncertainties in each C pool are generally proportional to the mean estimates. Variable importance (Figure 7) shows that prior data layers for each pool (e.g., 2010 AGB initial conditions, prior year’s annual mean LAI and SM, and SoilGrids SOC) are most important for explaining the current year’s SDA posterior mean. The 1-km gridded C estimates provide the potential for improved C conservation and management (e.g., optimize national C outcomes, Jia et al., 2025). Specifically, because most monitoring is done pool-by-pool, and indeed higher resolution maps do exist for most individual pools, C management is likewise usually done pool-by-pool. For example, one can currently register a C sequestration project with most voluntary markets that just considers management impacts on forest AGB without considering impacts on the overall ecosystem C budget (e.g., positive or negative impacts on SOC). This system also opens the door to more formal attempts to reconcile top-down and bottom-up C budgets, for example, by using our posteriors as land priors on atmospheric inversions or coupled land-atmosphere DA systems (de Rosnay et al., 2022). The associated uncertainty estimates, along with the spatial covariances, are critical to such efforts and could further improve the accuracy of MRV by accounting for the spatial correlations between locations within NA (MacBean et al., 2022).

The temporal trend map (Figure 8) shows that AGB is increasing in the Alaskan tundra, decreasing across western US coast forests, and exhibiting mixed responses across the eastern CONUS. The increase in AGB in Alaskan regions shows similar trends to those reported by Yang et al. (2023), who found increased forest biomass over time, while the declining trend in AGB across the western coast forest is likely due to increases in wildfires (Jiang et al., 2025). While the temporal patterns of AGB across the eastern CONUS are more diffuse, this might be due to a discontinuity in the LandTrendr AGB after 2017 (Figure S12), which shows that AGB estimates across the 8,000 locations are, on average, lower in 2018 and 2019 than in 2017. LAI shows increasing trends in the northern Arctic regions, similar to the AGB increment, and the central US trend (attributed mostly to the increment towards agricultural regions), with decreasing trends across Canadian boreal forests. The temporal patterns of leaf carbon are important indicators of potential changes in forest ecosystems. The browning trend aligned well with the 2023 wildfire event across Canada (Byrne et al., 2024). The browning could be linked to forest disturbances such as wildfire (Shi et al., 2025), urban development, forest disease (Fahey et al, 2022), etc., which could be used to inform future forest field validation and management (Potapov et al., 2025). The SM exhibits a slight decline across eastern NA and an increase in tropical regions. Changes in soil moisture might be linked to climate change (Qin et al., 2023), which has shown a decline in global soil moisture. As a result, plants are more likely to be stressed by water limitations, thereby altering micro-to-global climate feedbacks (Fu et al., 2024). SOC remains mostly stable, with a slight rise in Alaska.

The uncertainty trends for each pool (Figure 9) indicate that AGB uncertainty generally increases. The increase in AGB uncertainty outside CONUS US is mainly due to the absence of direct data constraints, in which case AGB is only indirectly constrained by the model and the other three assimilated pools (Zhang et al., 2025). The increase in AGB uncertainty within CONUS US is likely due to a 2017 discontinuity in LandTrendr, which introduces additional data error and thus increased uncertainty, and the absence of a LandTrendr constraint in 2024. The LAI uncertainty shows a slight but consistent decline over northern forests and an increasing trend across the central US grasslands. The decreasing trend towards northern forests might be attributed to the decline in the pool size and the relative smaller phenological magnitude across the boreal conifers, compared to the central US areas, which are dominated by croplands. SM uncertainties are mostly stable, and the few areas where uncertainty increased over time are driven by SMAP itself. Overall, SOC uncertainty declines, with a greater reduction observed at higher latitudes.

### 4.2. What are the patterns, across space, time, and C pools, of the residual errors in the hybrid SDA when compared to the corresponding data constraints and other data products during the validation?

All four variables show high R^2^ and low RMSE compared to the associated data constraints (Figure 10). LAI and SM show consistent high accuracy over time, while AGB and SOC exhibit an initial “burn-in” period as the SDA nudges the model simulation towards the observations. The high accuracy for LAI and SM is likely due to the relatively small data uncertainty, which allows the data to dominate the assimilation. This observation is consistent with the observation that the analysis uncertainties were only slightly reduced relative to the observations. The small contribution of the process model to both LAI and SM posteriors is likely due to the fact that both vary substantially over time, whereas we only assimilated them annually, meaning that the previous year’s posterior provided minimal constraint on this year’s prediction (Zhang et al., 2025). Many studies have been conducted on monthly-to-seasonal DA for both LAI and SM that showed stronger contributions of the process model (Tian et al., 2022 & Zhuang et al., 2024), while Wang et al. (2024) acknowledge the critical role of high-frequency DA in agricultural applications.

In Figure 11, there is no large-scale systematic bias for any C and water budgets predictions, demonstrating the success of our ML emulator. AGB residuals (-10 to 10 kgC/m^2^) are more variable than SOC residuals (-4 to 4 kgC/m^2^), though in both cases larger residuals are associated with larger pool sizes (e.g., eastern and western CONUS forests and northern arctic SOC). In addition, the SOC residuals in northwestern NA are likely due to insufficient ISCN records (Table S1), introducing biases towards arctic ecosystems. While we achieved high accuracy for LAI and SM (Figure 10) and the residuals are expected to be spatially randomly distributed, there are regions where residuals are clustered (e.g., higher LAI in Alaska and lower LAI in eastern and western coastal regions; higher SM in arctic tundra and lower SM in northern forest). Those patches might be associated with a need to better account for non-stationarity within the ML emulator (Rollinson et al., 2021), which might lead to additional uncertainties in specific ecosystems.

Overall, we achieved high accuracy for the AGB validations against GEDI + ICESat-2 (R^2^ = 0.73), except for underestimated forests in the eastern and southern NA, which might be due to the saturation of LandTrendr products (spectral indices are not sensitive to changes in biomass, Zhao et al., 2016; Sa et al., 2024), as well as a need to improve the process model’s calibration for such high-biomass systems (Dokoohaki et al., 2022; Fer et al., 2018, 2021). The FIA BIGMAP validations show high accuracy across forested locations, with slight underestimations of AGB across deciduous forests (Figure 15). The SOC validations against ISCN observations show that SOC is generally well predicted, with most ecoregion comparisons falling within the ensemble confidence interval, except for ecoregions with limited ISCN records (Figure 16).

### 4.3. Can the ML bias-correction improve the accuracy of our predictions? What factors are associated with model biases across space, time, and the C pool?

By comparing SDA accuracy before and after applying the debiasing algorithm (Figures 10 and S2), we see that SOC shows the greatest improvement in accuracy (RMSE reduced from 4.9 to 1.8 kgC/m^2^), likely due to the static SOC assimilated into the workflow, which makes it easier to predict residuals in the last step. While the accuracy of AGB without the debiasing gradually increases and decreases across time, applying the debiasing leads to fast-converged and high accuracy (RMSE reduced from 1.63 to 0.79 kgC/m^2^) results relative to those without debiasing. The differences before and after applying the debiasing module are negligible for LAI and SM, likely because LAI and SM are more variable over time, making it challenging to predict residuals from the previous time step.

Comparisons of validation results before and after applying the debiasing algorithm suggest similar accuracy for the GEDI + ICESat-2 AGB across the CONUS scale, but with improved accuracy in northern and southern evergreen forests. The improvements toward the northern and southern NA are likely due to some of the constrained locations being split into the >50 and <30 latitude windows. The similarity in accuracy within CONUS before and after the debiasing suggests that the space of improvements is likely limited by the LandTrendr products, which are biased relative to GEDI estimates (Figure S10). As a result, the accuracy of AGB may not improve further relative to GEDI, even if the estimates improve relative to LandTrendr. There are also improvements in the accuracy of the FIA BIGMAP validations, with evergreen forests showing the largest gains. Finally, the ISCN validation shows that we achieved higher accuracy after applying the debiasing module, reducing the number of outliers from 6 to 3. The variable importance analysis (Figure 17) shows that current SIPNET forecasts, residual lag, and the associated data layers mainly explain the residual errors of four state variables (i.e., the process model’s biases are more driven by autocorrelation than specific environmental conditions).

### 4.4 Future Directions

A top priority for improving the SDA would be to increase the number of data constraints. Assimilating AGB estimates from LiDAR platforms (e.g., ICESat-2 and GEDI) would expand the spatial coverage of AGB estimates and mitigate saturation in optical remote sensing. In addition, disturbances are known to have large impacts on stock changes, land-atmosphere fluxes, and C export from ecosystems (Williams et al., 2014; Kasischke et al., 2013), but were not included in the SIPNET model itself. Although the land cover and disturbance history were included in the ML-downscaling algorithm as covariates, disturbance impacts are likely underestimated. For example, in this study, we didn’t observe the significant lose of woody biomass given the browning trend across Canada. This could be addressed by including disturbance events in the SDA (Dietze et al preprint), leveraging information about disturbance in both the SDA and through additional disturbance data constraints (Pickens et al., 2025). Given the limitations of ISCN records, initializing SOC using the Northern Circumpolar Soil Carbon Database (NCSCDv2, Hugelius et al., 2013) could reduce uncertainty in our SDA workflow.

To address the biases identified by ML debiasing, an important future direction is to refine the model calibration to include more plant functional types or to account for ecological variation in model parameters across space, as is done in calibration-only systems like CARDAMOM (Yang et al., 2022). In the longer term, it would be valuable to integrate the SDA and calibration, so that data constraints are being used to update both model states and parameters simultaneously, which should lead to more accurate C cycle estimates than either SDA or calibration alone. Beyond that, to better account for the temporal variabilities of LAI and SM, a seasonal-to-monthly SDA would be valuable to improve model forecasts and error predictions during the debiasing step.

While the predicted maps align well with existing knowledge of the four C and water pools across NA, the current ML emulator has some limitations. For example, it assumes independence across ensemble members, space, and time for each prediction, which may overestimate the uncertainties across space and time. A future direction would be to use a spatiotemporal ML emulator (e.g., spatio-temporal graph neural networks (STGNNs), Wen et al., 2023) to improve error estimates and allow the sub-annual temporal downscaling of fast-changing pools (e.g., LAI, SM) and fluxes. Also, the emulator doesn’t include ML uncertainty calculations (i.e., errors during training), which underestimates the uncertainty in the simulated maps. Bayesian Neural Network (BNN, Saad et al., 2024) or ensemble ML predictions (i.e., utilizing different ensemble ML methods to represent uncertainty, Ganaie et al., 2022) may improve the propagation of these uncertainties.

## 5. Conclusions

Compared to a status quo dominated by coarse-resolution (0.5°) modeling without direct data constraints, this study demonstrates the feasibility of estimating land C budgets at a much higher resolution (1 km) across continental scales by harmonizing remotely sensed observations, ground-based data, and process-based ecological modeling. The SDA analysis and gridded estimations provide up-to-date NA C budgets aimed at accurate C MRV. Our validation analyses show relatively high agreement with both in-sample and held-out data sources, as well as reduced uncertainty compared to the data constraints. In the future, we anticipate that SDA will become a core technology for providing accurate and up-to-date C MRV, continuously constrained by multidimensional observations from around the world.

