## Supplements 1-13 for "Mapping the North American Terrestrial Carbon Cycle: A Process-based Reanalysis Using State Data Assimilation (SDA)"

**Supplementary Tables.**

*Table S1. Information on the aggregated ISCN database across the level-2 EPA ecoregions. Here, we show the ecoregion ID, ecoregion name, number of ISCN records, and the averaged SOC density.*

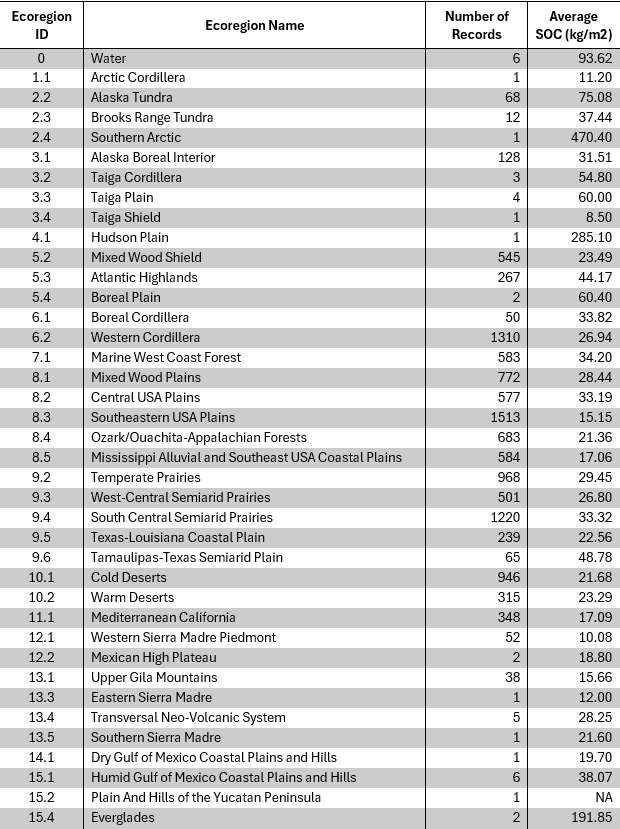

**Supplement 1. The workflow of generations for soil water holding capacity and drainage rate.**

The soil moisture used in the model is limited by the soil water holding capacity. This critical parameter and drainage rate were estimated through a multi-step process:

- Soil Texture Data: We used the mean and quantiles (5th, 50th, and 95th) of sand, clay, and silt fraction data from SoilGrids (Hengl et al., 2017) for each site and depth.
- Ensemble Generation: These data were used to fit Dirichlet distributions using the moment matching method, which fit the quantiles with means as known moments. Then, the 100-ensemble soil texture datasets were generated for each site and depth.
- Parameter Inference: Soil physical properties, including water holding capacity and drainage rate, were inferred for each ensemble member using pedotransfer equations (Cosby et al., 1984).
- Integration: The final soil parameters were integrated over the top 100 cm to align with the depth of the SMAP soil moisture data for each site.

**Supplement 2. The workflow of machine-learning (ML) LandTrendr AGB uncertainty predictions.**

As mentioned in the main text, the LandTrendr AGB we used provides mean AGB estimates from 2012 to 2023, whereas the uncertainty estimates end in 2017. Therefore, we used the ML method to train and predict uncertainties from 2017 onward. Here are the workflows that were used to predict the uncertainties:

- We extracted spatial attributes for the selected locations within CONUS from Table 2 in the main text, including annual mean ERA5 climate, DEM, stand age, MODIS land cover, and soil physical properties. Note that we replaced the 2010 global AGB layer with the corresponding LandTrendr AGB mean estimates for each site and year.
- After that, we train our random forest model (Y ~ X, where X represents the stacked spatial attributes discussed above, and Y is the LandTrendr AGB uncertainty estimates for the corresponding location and time).
- Then we predict AGB uncertainties using the same attributes X (within which the LandTrendr AGB mean layer should fall between 2018 and 2023).

**Supplement 3. Relationship between ORC (organic carbon content) and OCD (organic carbon density).**

OCD [kg/m3] = ORC [%]/100 × BLD [kg/m3] × (1-CRF[%]/100), where BLD is the bulk density, CRF is the coarse content fraction.

**Supplement 4. Differences in DOY (day-of-year) between the date of selected MODIS LAI observations and the target data assimilation dates (July 15th each year from 2012 to 2024).**

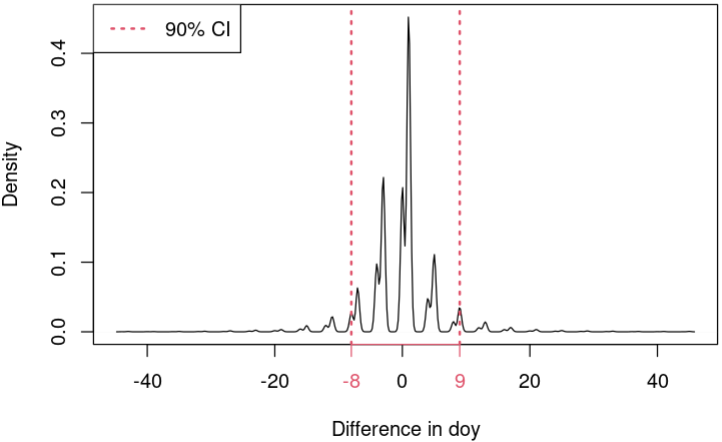

*Figure S1. The differences in the day-of-year of our LAI selection date relative to July 15th for each year and site.*

**Supplement 5. SIPNET Model.**

We used the Simplified Photosynthesis and Evapotranspiration Model (SIPNET) (https://github.com/PecanProject/sipnet), a simple pool-based process model, as the scaffold for making forward predictions of all C pools and fluxes and reconciling predictions with observations (Braswell et al., 2005). SIPNET was selected based on its low computational cost and its simplified process representations of coupled carbon and water cycling, which facilitate interpretability and data assimilation.

SIPNET contains four vegetation carbon pools (leaf, stem, coarse roots, and fine roots) and one soil carbon pool (Figure 5). Photosynthesis is calculated using a light-use-efficiency approach that is a function of maximum photosynthesis, LAI, and photosynthetically active radiation, with modifiers for air temperature, vapor pressure deficit, and soil moisture. SIPNET also has a simple representation of vegetation C allocation based on fixed fractions to each plant carbon pool. For deciduous systems, SIPNET was run in “prescribed phenology” mode based on the day-of-year of leaf-on and leaf-off. Losses from plant carbon pools are based on fixed turnover rates and maintenance respiration, which is a Q10 function of temperature. Soil carbon is increased by wood and leaf litter and reduced by soil respiration. Finally, soil moisture is modeled using a simple bucket model approach, with precipitation as input and evaporation, transportation, and drainage as outputs. To match the assimilated observations, SOC was modeled down to a 2m depth, while soil moisture was modeled to a 1m depth.

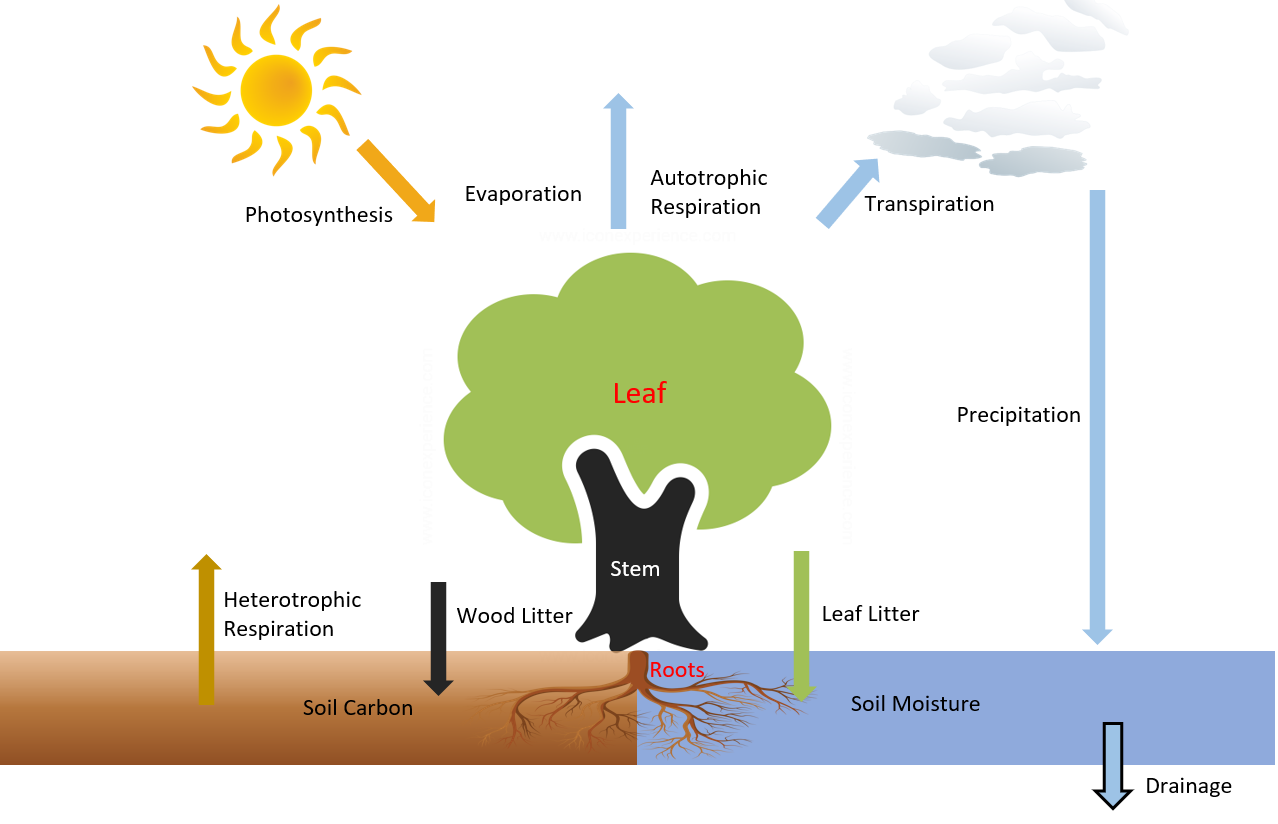

*Figure S2. Flowchart of the SIPNET model. The model includes four vegetation carbon pools (leaf, stem, coarse roots, fine roots) and one soil carbon pool. Photosynthesis adds carbon to the system, which is then allocated to the four vegetation pools. Meanwhile, autotrophic and heterotrophic respiration remove carbon from the vegetation and soil, respectively. Vegetation pool turnover (a.k.a. litter) adds to soil carbon, and precipitation adds soil moisture, the latter of which is removed by transpiration, evaporation, and drainage.*

**Supplement 6. Machine Learning Debias workflow.**

Systematic errors in model forecasts can persist across cycles and reduce the skill of the data assimilation. To address this, we developed a machine learning (ML) framework that corrects ensemble forecasts before they are passed into the filter. The goal is to shift the ensemble mean to remove bias while leaving the ensemble spread intact, so that uncertainty propagation remains governed by the filter.

At each assimilation cycle, the process-based model (SIPNET) generates an ensemble of forecasts for each state variable. From this ensemble, we compute the mean forecast, denoted μᶠ, and the ensemble covariance, denoted Σᶠ. Given an observation yᵒᵇˢ, we define the residual as ε = yᵒᵇˢ − μᶠ. These residuals are then merged with a set of environmental covariates, including climate, soils, land cover, disturbance history, and topography. In addition, the forecast itself was explicitly included as a predictor, since systematic errors often scale with forecast magnitude.

The debias model is a stacked regression framework implemented in Python and called from R via the reticulate interface. For each state variable, we fit two base learners: a K-Nearest Neighbors regressor (with the number of neighbors k selected by cross-validation) and an ExtraTrees ensemble. Both models are trained to map the combined predictor set (covariates plus forecast) to the observed residuals. Their predictions are then blended with a weight w between 0 and 1, chosen to minimize root mean squared error on the training set. The resulting blended model produces a predicted residual ε̂ for the next timestep.

To apply the correction, we add the same offset ε̂ to each ensemble member. In other words, the corrected ensemble is given by yᶠ,ᶜᵒʳʳ = yᶠ + ε̂. This procedure shifts the ensemble mean (μᶠ,ᶜᵒʳʳ = μᶠ + ε̂) while leaving the covariance unchanged (Σᶠ,ᶜᵒʳʳ = Σᶠ). By construction, the debiasing step only adjusts the location of the ensemble distribution, ensuring that spread and shape are preserved for the subsequent assimilation update.

These bias-reduced forecasts are then passed to the Tobit-Gamma Ensemble Filter (TGEnF) for assimilation with observations. Cross-validation confirmed that including the forecast itself as a predictor significantly improved residual prediction, and feature importance analysis showed that the forecast was consistently the single most important predictor across all variables. Environmental covariates such as disturbance history, precipitation, temperature, and soil texture provided additional explanatory power, but their effects were secondary to the forecast itself. Together, these results show that the ML debiasing framework reduces systematic forecast–observation mismatches and stabilizes the assimilation process by aligning the prior mean with observational evidence before the filter update.

**Supplement 7. Differences in performance between XGBoost and Random Forest regression models.**

In this study, we first assessed the performance of the Random Forest and XGBoost machine learning models to determine which method to use for interpolating SDA outputs. After completing the NA SDA experiments for the preselected 8,000 locations and four variables, we randomly selected 50% of the samples from the SDA analysis for each year and variable, across each MODIS landcover class, to avoid oversampling specific land covers. Then we trained both the Random Forest and XGBoost models on 50% of the samples and predicted the remaining locations. As shown in Figure S3, RandomForest consistently outperforms XGBoost across variables and time periods. Therefore, we will proceed with the RandomForest method as the NA SDA emulator. For the across-variable predictability, SOC is generally easier to predict, while LAI predictions are comparatively more difficult. From the temporal patterns, AGB, LAI, and SOC show increased accuracy over time, although LAI exhibits more interannual variability.

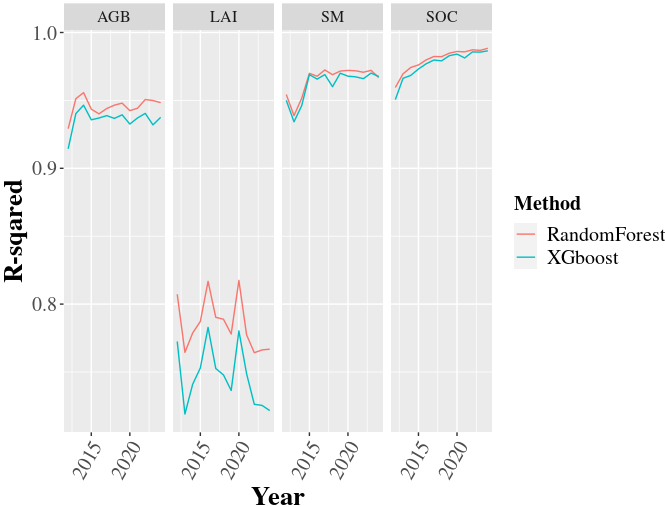

*Figure S3. The comparisons of R-squared of the out-of-sample prediction accuracies between the RandomForest and XGboost methods. Here, we first randomly sampled 50% of the 8,000 preselected locations across each land cover class and split the corresponding SDA analysis results as the training dataset. Then, we used both regression models to predict the remaining 50% of the locations and calculate the R-squared value between the model predictions and the SDA analysis results.*

**Supplement 8. Validations between our SDA results and the held-out datasets for AGB and SOC.**

Here, we ran SDA experiments without the debiasing module, and we first calculated the temporal trends of the accuracy across variables between SDA analysis and the corresponding data constraints (Figure S4), then validated the AGB against the GEDI+ICESat 2 (Figure S5) and FIA BIGMAP (Figure S6). We finally validated the SOC analysis against the ISCN database across the aggregated EPA level 2 ecoregions (Figure S7).

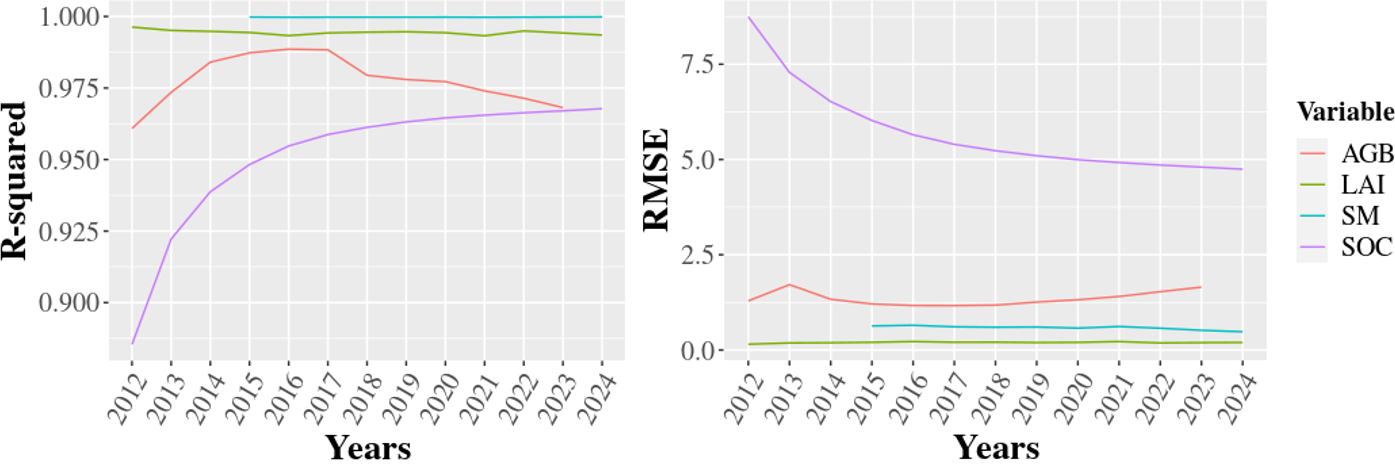

*Figure S4. The temporal trends (before debiasing) of different error metrics (R-squared and RMSE) from 2012 to 2024 across C and water budgets. Here, we compared our ensemble mean estimates against the corresponding measurements for the pre-selected 8,000 locations across time. Note that any missing values represent NAs in the observation (e.g., SMAP SM available after 2015).*

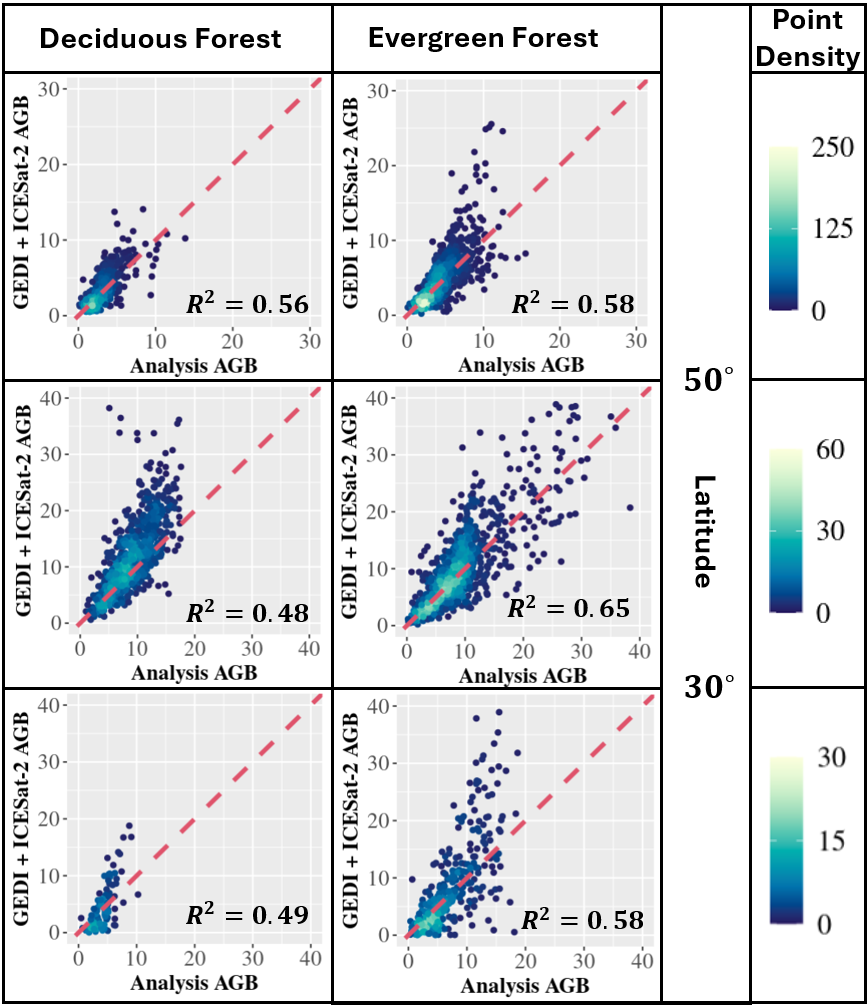

*Figure S5. The comparisons between GEDI + ICESat-2 and our analysis of AGB results before debiasing. Here, we split the comparisons into three rows by the corresponding latitude range (> 50 degrees,> 30 and < 50 degrees, < 30 degrees) and two columns by forest types (deciduous and evergreen forest, respectively). The point density colors the dots: yellower means more points are clustered, while bluer means fewer points are clustered.*

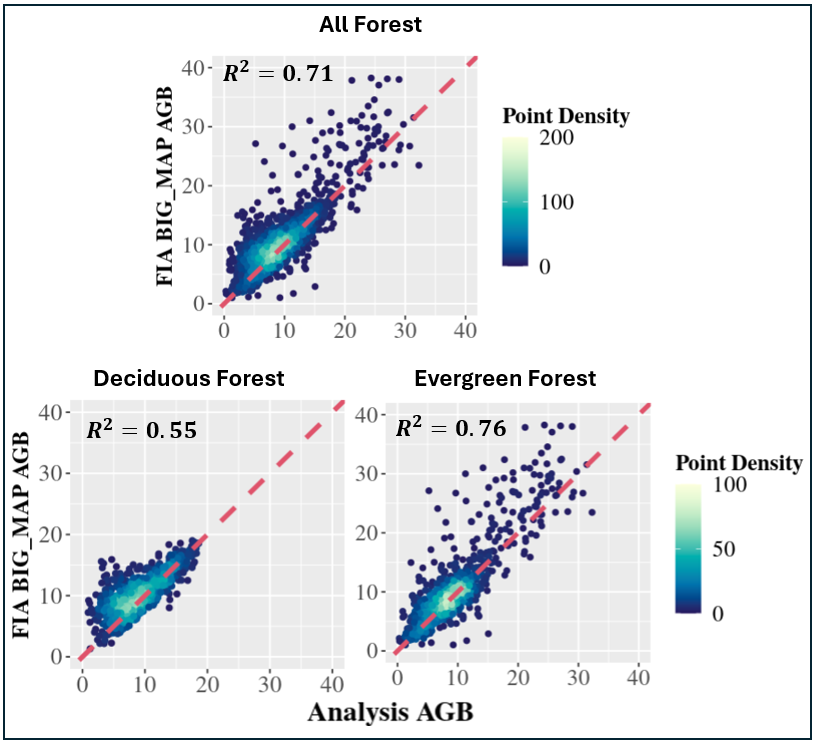

*Figure S6. Comparisons between the AGB analysis (averaged from 2014 to 2018) of preselected locations and the FIA BIGMAP database across (A) all forest, (B) deciduous forest, and (C) evergreen forest before debiasing.*

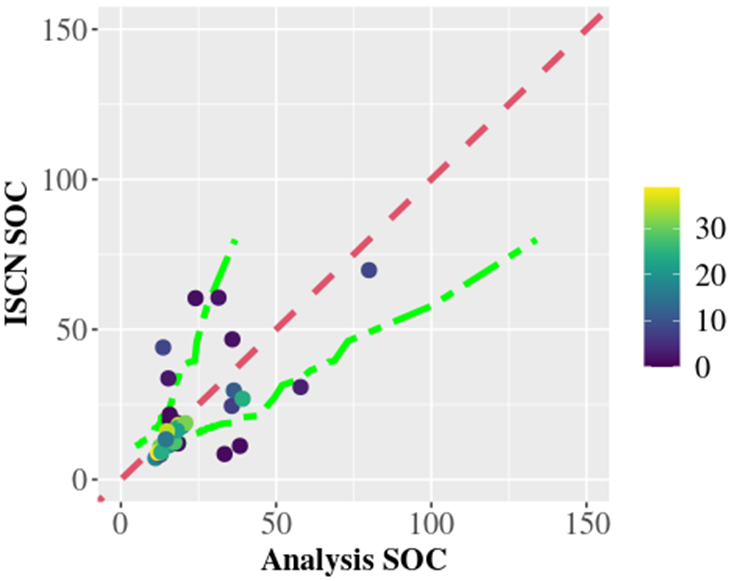

*Figure S7. Comparisons between ISCN and the Analysis of SOC before the debiasing. Here, we first calculate the aggregated ISCN SOC mean values across the level 2 EPA ecoregions. After that, for each corresponding ecoregion, we calculate the quantiles (2.5, 50, and 97.5%) from the ensemble analysis results. Finally, we plot the mean comparisons (dots colored by the square root of the number of available ISCN records for each ecoregion), one-to-one reference line (in red), and the upper and lower boundaries (in green) of the ensemble analysis results.*

**Supplement 9. The relationship between pool sizes and their uncertainties.**

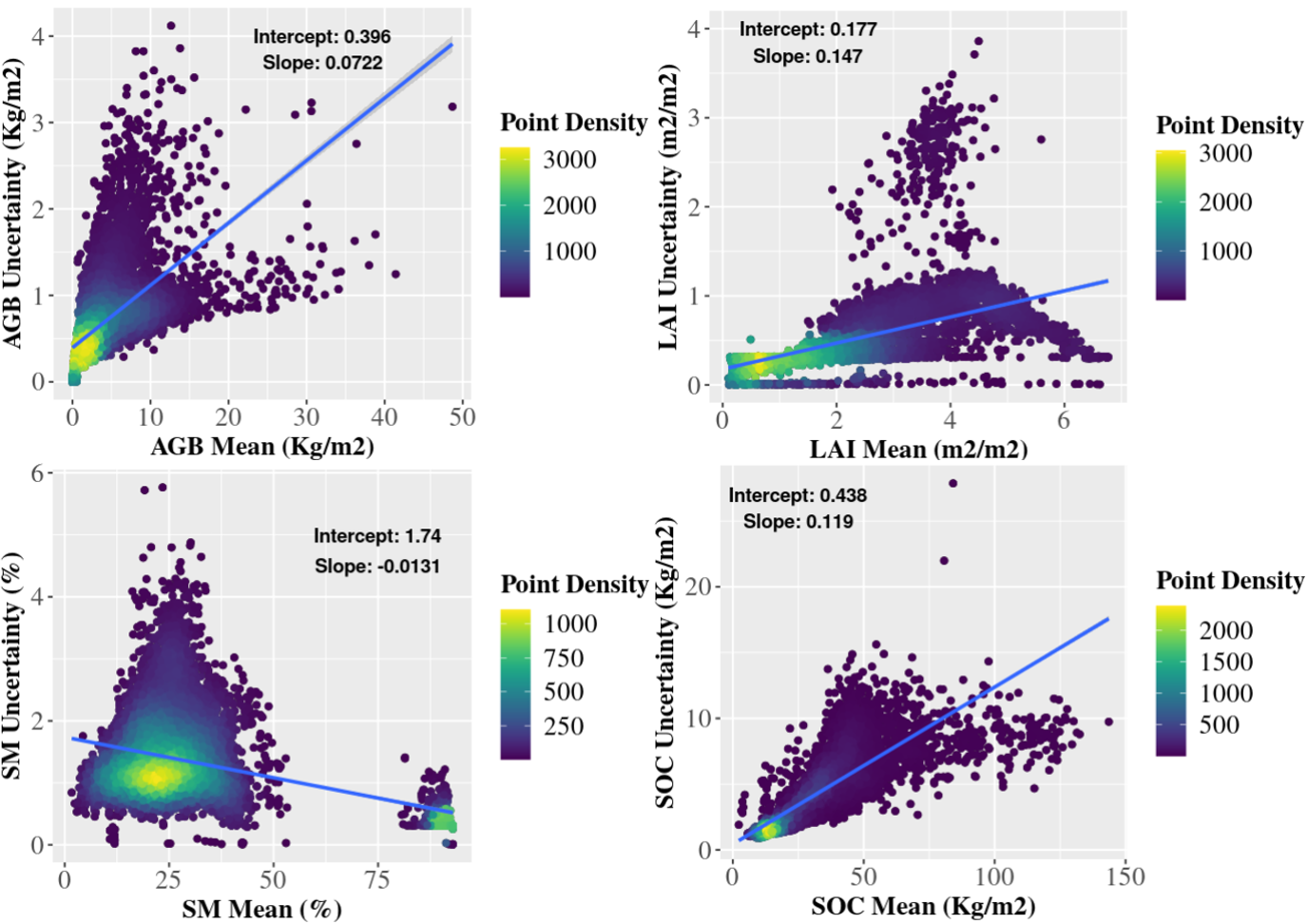

*Figure S8. Relationships between the mean and uncertainty across different C and water pools.*

**Supplement 10. Validation of AGB between SDA analysis and GEDI+ICESat-2 AGB estimates from 2019 to 2023.**

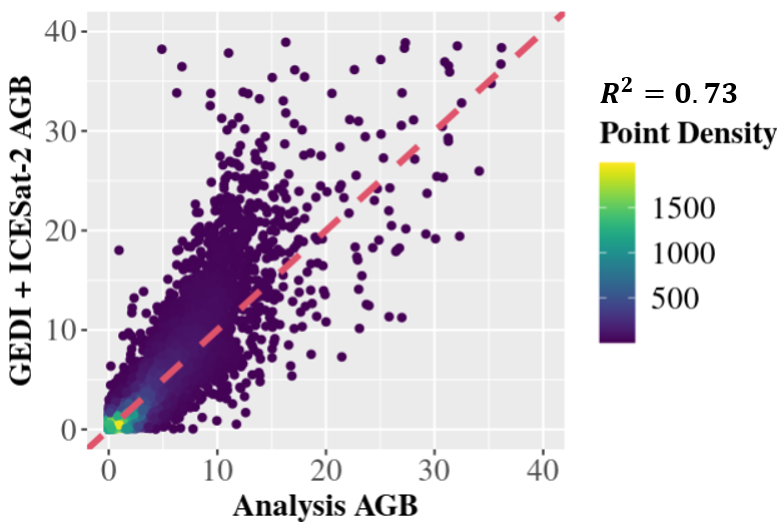

*Figure S9. Comparisons of AGB between our analysis results and the ICESat-2 and GEDI products.*

**Supplement 11. Comparisons between LandTrendr and GEDI AGB estimates across forest types.**

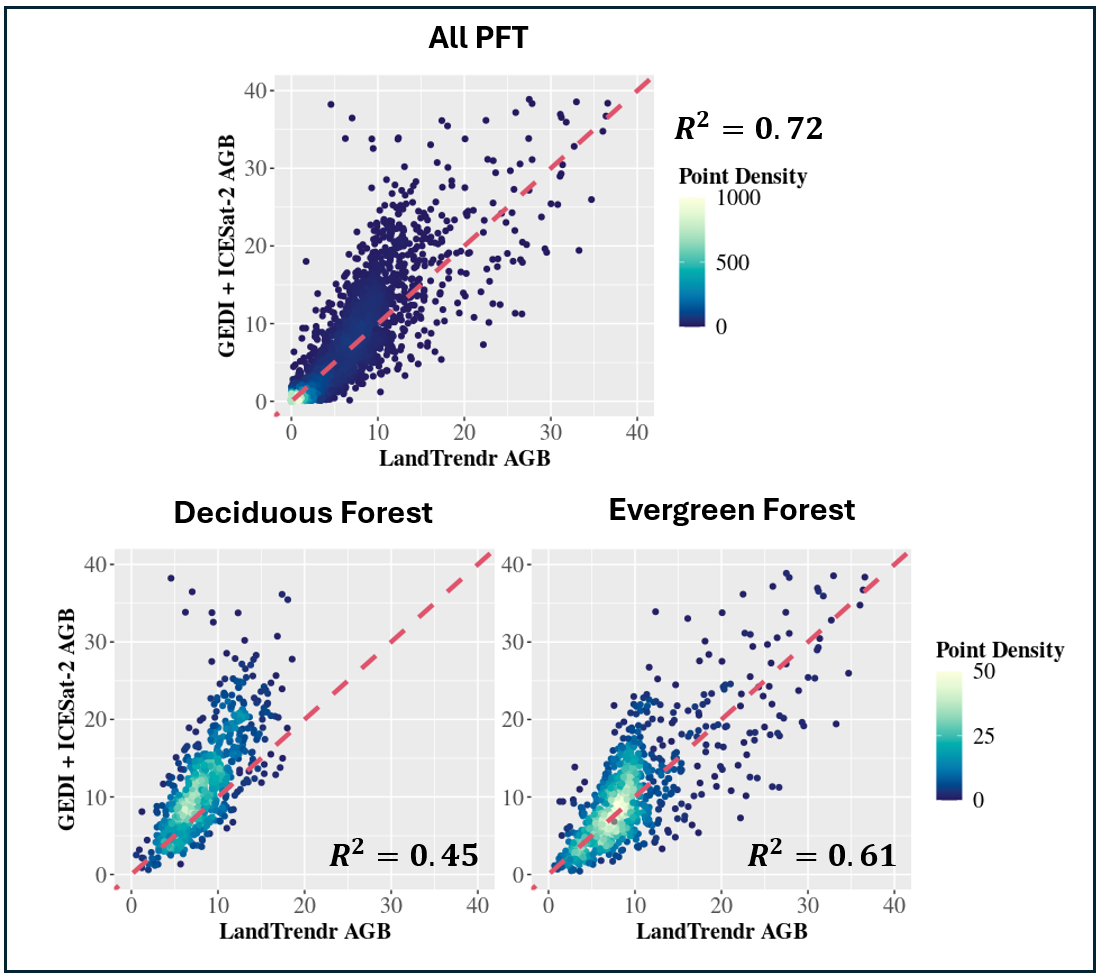

*Figure S10. Comparisons between LandTrendr and ICESat-2 + GEDI AGB estimations for 1) All PFTs; 2) Deciduous; and 3) evergreen forests within the CONUS US scale.*

**Supplement 12. The residual error map of AGB between SDA outcomes and FIA BIGMAP.**

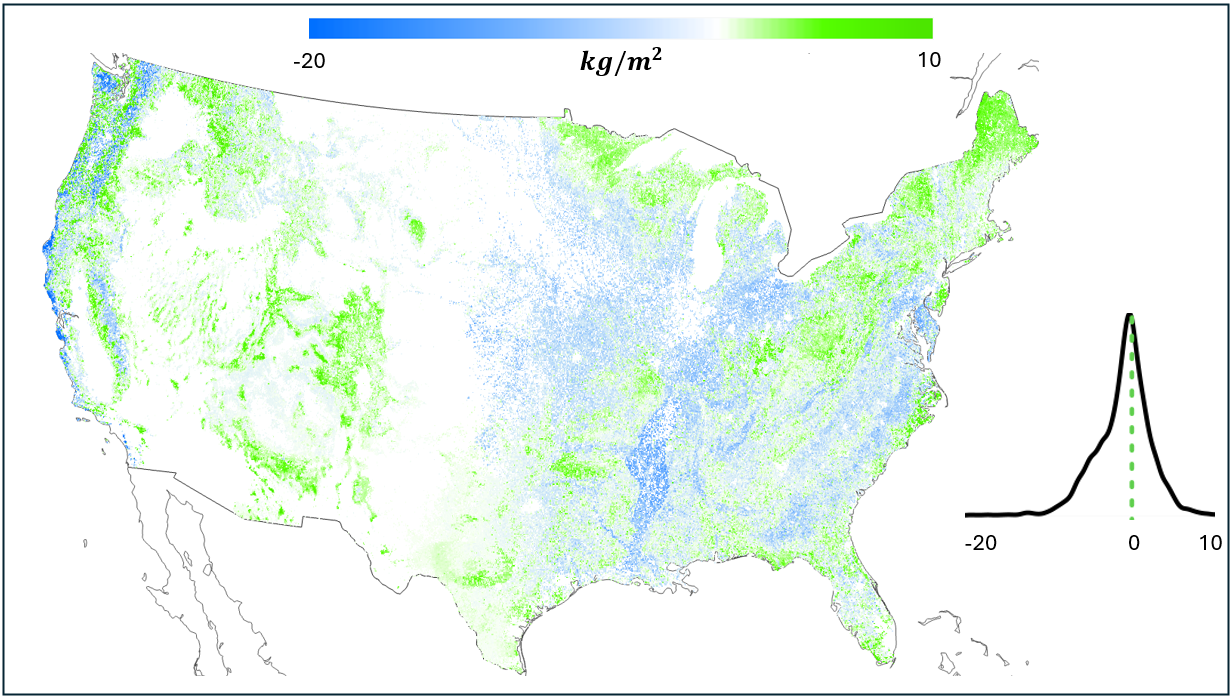

*Figure S11. Map of residual error between the SDA AGB Analysis mean (averaged from 2014 to 2018) and the FIA BIGMAP.*

**Supplement 13. The discontinuity of the LandTrendr AGB estimates at year 2017 across 8,000 locations.**

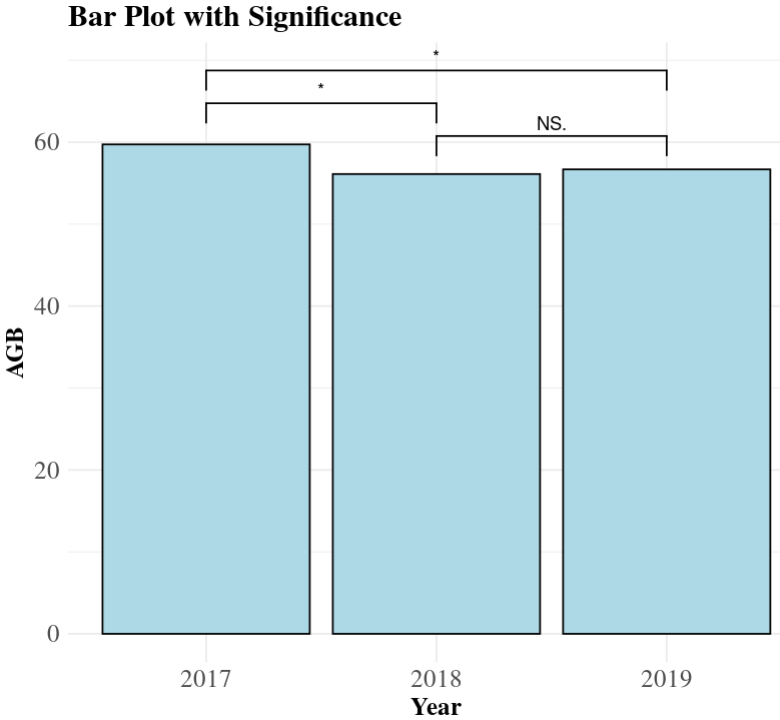

*Figure S12. Barplot of LandTrendr AGB observations across selected 8,000 locations from 2017 to 2019. From this figure, we can see that the LandTrendr AGB is statistically significantly reduced after 2017.*
